# IMD-mediated innate immune priming increases Drosophila survival and reduces pathogen transmission

**DOI:** 10.1101/2023.02.22.529244

**Authors:** Arun Prakash, Florence Fenner, Biswajit Shit, Tiina S. Salminen, Katy M. Monteith, Imroze Khan, Pedro F. Vale

## Abstract

Invertebrates lack the immune machinery underlying vertebrate-like acquired immunity. However, in many insects past infection by the same pathogen can ‘prime’ the immune response, resulting in improved survival upon reinfection. Here, we investigated the generality, specificity and mechanistic basis of innate immune priming in the fruit fly *Drosophila melanogaster* when infected with the gram-negative bacterial pathogen *Providencia rettgeri*. We find that priming in response to *P. rettgeri* infection is a long-lasting and pathogen-specific response. We further explore the epidemiological consequences of immune priming and find it has the potential to curtail pathogen transmission by reducing pathogen shedding and spread. The enhanced survival of individuals previously exposed to a non-lethal bacterial inoculum coincided with a transient decrease in bacterial loads, and we provide strong evidence that the effect of priming requires the IMD-responsive antimicrobial-peptide *Diptericin-B* in the fat body. Further, we show that while *Diptericin B* is the main effector of bacterial clearance, it is not sufficient for immune priming, which requires regulation of IMD by peptidoglycan recognition proteins. This work underscores the plasticity and complexity of invertebrate responses to infection, providing novel experimental evidence for the effects of innate immune priming on population-level epidemiological outcomes.

## Introduction

Immunisation using attenuated or inactivated pathogens is one of the most successful public health practices to reduce the incidence of infectious diseases(Pollard and Bijker, 2021). Immunisation works because humans and other vertebrate animals have evolved an acquired immune response capable of specific immune memory, which ensures a strong, precise, and effective response to a secondary infection(Boehm and Swann, 2014). Insects possess a robust innate immune response to pathogens which includes both cellular and humoral components (Hoffmann, 2003; Hultmark, 2003; Buchon et al., 2014), but lack vertebrate-like specialized immune cells responsible for acquired immunity. These differences in immune physiology resulted in the long-standing assumption that invertebrates should not be capable of immune ‘memory’, though this view was clearly at odds with empirical evidence from several invertebrate host-pathogen systems(Lanz-Mendoza et al., 2024; Milutinović and Kurtz, 2016; Prakash and Khan, 2022). There is now substantial evidence that insects also possess a form of “immune priming”, where low doses of an infectious pathogen, or even an inactivated pathogen, can lead to increased survival upon reinfection (Kurtz and Franz, 2003; Sadd and Schmid-Hempel, 2006; Lanz-Mendoza et al., 2024; Contreras-Garduño et al., 2016; Milutinović and Kurtz, 2016).

There are many ways in which invertebrates may enhance their immune responses upon reinfection(Arch et al., 2022; Gomes et al., 2022; Prakash and Khan, 2022). In *Drosophila,* the priming response during infection with the gram-positive bacterial pathogen *Streptococcus pneumoniae* was shown to be dependent on haemocytes and phagocytosis, while the Toll-pathway - the main pathway involved in clearance of gram-positive bacteria - was shown to be insufficient for successful priming (Pham et al., 2007). Increased phagocytic activity in primed individuals has also been shown to play a key role in priming during *Pseudomonas aeruginosa* infection in Drosophila(Apidianakis et al., 2005), while in the woodlouse, prior exposure to heat-killed bacteria led to increased phagocytosis by haemocytes upon reinfection (Roth and Kurtz, 2009). In other *Drosophila* work, *PGRP-LB* (peptidoglycan recognition protein *LB*, a negative regulator of the Immune deficiency (IMD)- pathway) was identified as a key mediator of transgenerational immune priming against infection with parasitoid wasps (*Leptopilina heterotoma* and *Leptopilina victoriae*). Here, downregulation of *PGRP-LB* was necessary to increase haemocyte proliferation, required for wasp encapsulation by haemocytes in the offspring (Bozler et al., 2020). By contrast, in response to infection with *Drosophila C virus* (DCV)*, Drosophila* progeny can produce a DCV-specific priming response by inheriting viral cDNA from the infected adult flies which is a partial copy of the virus genome (Mondotte et al., 2018, 2020). The benefits of priming are not always associated with increased pathogen clearance, as shown during Enterococcus faecalis infection, where protection was explained by increases tolerance of infection, not increased clearance (Cabrera et al., 2023).

Beyond its underlying mechanisms, an important but understudied aspect of immune priming is its potential consequences for pathogen transmission (Gomes et al., 2022; Tate, 2017, 2016; Tidbury et al., 2012). Epidemiological modelling predicts that priming should affect the likelihood of pathogen persistence, destabilise host–pathogen population dynamics, and that these effects depend on the degree of protection conferred by priming (Tidbury et al., 2012). Further theoretical work suggests that priming can either increase or decrease infection prevalence depending on the extent to which it affects the pathogen’s colonization success and the host’s ability to clear or tolerate the infection (Tate, 2017). While primed individuals may live longer, thus extending the infectious period, their pathogen burden may be lower, which could lead to lower pathogen shedding and less severe epidemics. Immune priming is therefore expected to have a significant impact on the outcome of pathogen transmission by directly modifying important epidemiological parameters, but the strength and direction of these effects is not intuitive to predict.

Here, we focus on immune priming in *Drosophila* when infected with the gram-negative bacterial pathogen *Providencia rettgeri* to investigate the occurrence, generality, duration, and mechanistic basis of immune priming during systemic infection. Furthermore, motivated by the theoretical predictions about the role of immune priming on epidemiological dynamics, we also designed transmission experiments to enable us to test how immune priming could affect each of these behavioural and immunological components of pathogen spread.

## Results

### Immune priming in *Drosophila* is a long-lasting response, showing sex-specific effects in different genetic backgrounds

We first examined if the length of time between the initial non-lethal exposure with heat-killed *P. rettgeri* and the secondary pathogenic challenge with live *P. rettgeri* affects the extent of priming. To address this, we exposed *w^1118^* male and female control flies to live *P. rettgeri* 18-hours, 48-hours, 96-hours, 1-week or 2-weeks following the initial exposure to heat-killed bacteria. Male *w^1118^* flies showed increased survival after initial priming for time points 18-hours, 48-hours and 96-hours, and still showed a significant, albeit reduced, priming response 1-week and 2-weeks after the initial exposure (**Fig**. 1 A-E). Female flies did not show a priming response when infected 18-hours following the initial challenge, but the priming response increased with 48-hours and 96-hours priming intervals before completely disappearing after a week time-interval (including 2-weeks) (**Fig**. 1A-E, **Table**-S1).

**Figure 1:**
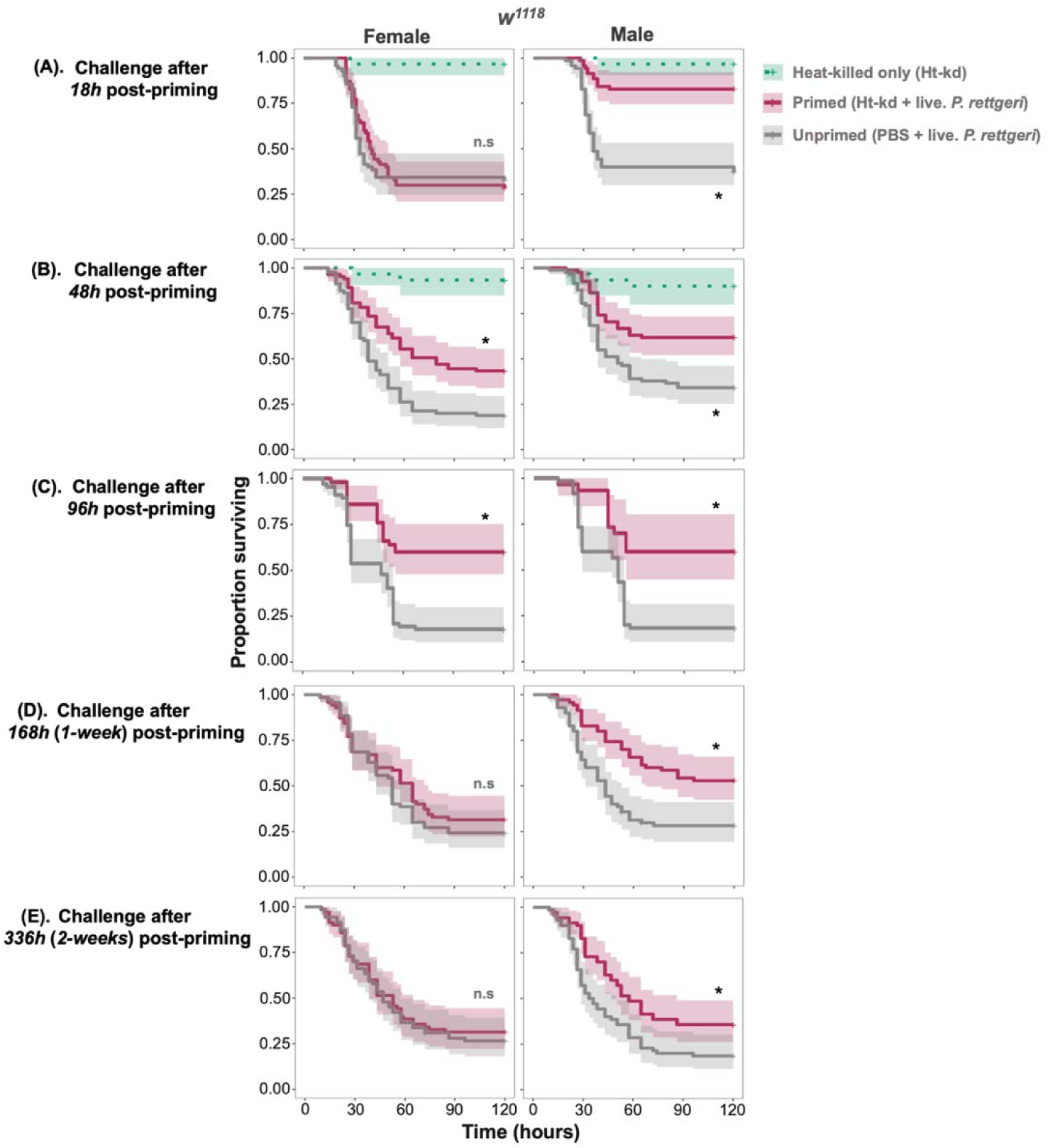
The benefit of priming is long-lasting. Survival curves of *w^1118^* females and males with primed (exposed to heat-killed *P. rettgeri* in the first exposure) and unprimed (exposed to a sterile solution in the first exposure) treatments challenged with live *P. rettgeri* pathogen after (A) 18-hours (B) 48-hours and (C) 96-hours (D) 168-hours/1-week and (E) 336-hours/2-weeks post priming that is, initial non-lethal exposure to heat-killed *P. rettgeri* (n=7-9 vial with 10-15 flies in each vial/sex/treatment/timepoint).

Next, we asked if priming occurred in two other commonly used *Drosophila* lines, Canton-S and Oregon-R (Ore-R) as observed with *w^1118^*. Given the widespread effects of the endosymbiont *Wolbachia* on *Drosophila* immunity (Brownlie and Johnson, 2009; Teixeira et al., 2008; Vanika Gupta et al., 2017) we also tested whether the presence of *Wolbachia* had any effect on immune priming by comparing the priming response of Oregon-R (OreR), the line originally infected with *Wolbachia* strain wMel, henceforth OreR^Wol+^ and a *Wolbachia*-free line OreR^Wol-^ that was derived from OreR^Wol+^ by antibiotic treatment (Vanika Gupta et al., 2017). Females and males of the four *Drosophila* lines were treated first with heat-killed *P. rettgeri*, followed by infection with live *P. rettgeri* 96-hours after the first treatment. Since the 96-hour time gap between priming and live *P. rettgeri* treatments showed maximum priming response (difference in survival) for both males and females of the *w^1118^*line (Fig. 1C), we kept the 96-hours timepoint as the consistent time-gap between priming and live infection throughout the study (**Fig**. 2C). Canton*-*S flies showed increased survival following priming and we observed this survival benefit across both sexes, although stronger in females (**Fig**. 2B, **Table** S2B). Oregon-R males also exhibited increased survival following priming, but priming had no significant effect on the females (**Fig**. 2C, **Table**-S2). We found that the presence of *Wolbachia* significantly improved overall survival of both males and females (**Fig**. 2C and D, **Table**-S3). However, the immune priming observed in males in absence of *Wolbachia* (OreR^Wol-^) was no longer present in flies carrying *Wolbachia* (OreR^Wol+^) (**Fig.** 2D, **Table**-S3).

**Figure 2:**
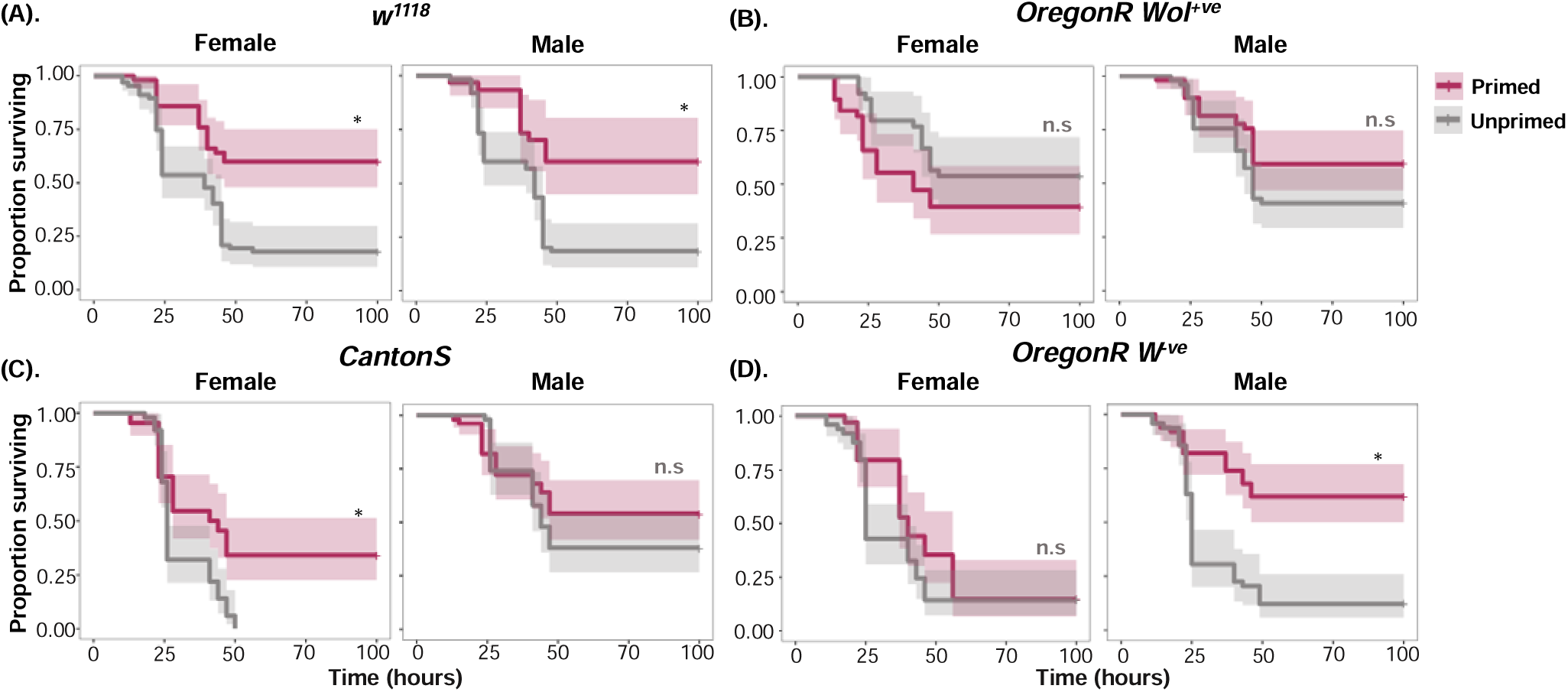
Survival curves of females and males of the commonly used *Drosophila* lines either primed-challenged or unprimed-challenged with *P. rettgeri*. (A) w^1118^ (B) Canton-S (C) Oregon-R carrying endosymbiont *Wolbachia* (D) Oregon-R cleared of *Wolbachia*. (n=7-9 vial with 10-15 flies in each vial/sex/treatment/line).

### The strength of immune priming is bacteria species-specific

A notable feature of immune priming is its specificity (Roth et al., 2009; Khan et al., 2017), which suggests an added level of functional complexity of insect immunity. We tested whether *D. melanogaster* also shows specific immune priming against *Providencia rettgeri (Pr)*. We used males of the *w^1118^* line, as males were shown to have stronger priming response than females and two pathogens, one closely related species of *P. rettgeri* i.e., *Providencia burhodogranariea (Pb)* and another less closely related gram-negative bacteria i.e., *Pseudomonas entomophila (Pe)*. Male flies primed and challenged homologously (*Pr – Pr*) species showed significantly improved survival rates compared with the heterologous treatment combinations (primed with *Pr* followed by challenge with other bacteria) and corresponding control (primed with PBS – challenge with live bacteria) (**Fig**. 3A-C, **Table** S4A-B).

**Figure 3:**
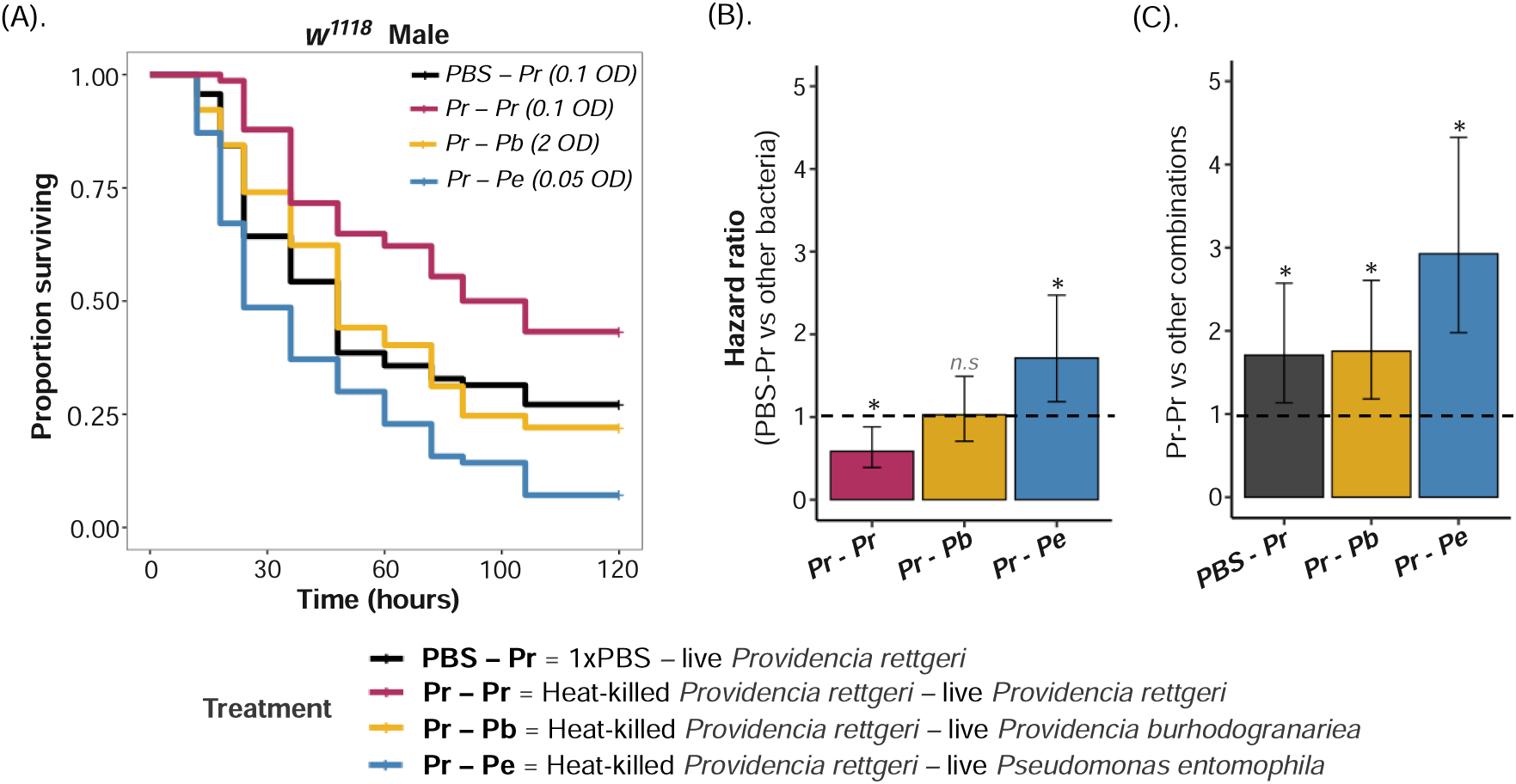
Bacterial species-specific immune priming response in *w^1118^* males. Two closely related species of Providencia bacteria *Providencia rettgeri* (Pr) and *Providencia burhodogranariea* (Pb) and another gram-negative bacteria *Pseudomonas entomophila* were used for testing the immune priming specificity in *Drosophila*. (A) survival curves (B) hazard ratio of unprimed males with 1x PBS followed by live *P. rettgeri* against males with primed *P. rettgeri* followed by live infection with other bacterial species *Pb* and *Pe*. (C). hazard ratio of primed males with heat-killed *P. rettgeri* followed by live *P. rettgeri* against males with primed *P. rettgeri* followed by live infection with other bacterial species *Pb* and *Pe*.

### Primed male *w^1118^* flies exhibit a transient reduction in bacterial load

Previous studies have shown that the priming response to fungi and gram-positive bacteria can be explained by increased clearance of bacterial pathogens in primed individuals (Pham et al., 2007; Khan et al., 2019). We therefore examined whether the increased survival following a prior challenge we observed was a result of greater bacterial clearance in the primed individuals, or if the primed flies were simply better able to tolerate the bacterial infection (Cabrera et al., 2023). To investigate this, we repeated the priming experiment with *w^1118^*as it showed the strongest priming phenotype in both sexes (**Fig**. 1C and **Fig**. 4A), and measured the bacterial load at 24-hours and 72-hours post-infection for both primed and unprimed female and male flies. Again, we observed a clear priming effect in both sexes (**Fig**. 4A, **Table**-S5). In primed males the systemic infection resulted in higher survival and also in decreased bacterial loads at 24-hours post-infection when compared to unprimed individuals (**Fig.** 4, **Table**-S5 for survival, **Table**-S6 for load). However, by 72-hours post-infection, bacterial loads had dropped in both primed and unprimed male flies and there was no detectable effect of priming on the bacterial loads (**Fig**. 4B, **Table**-S6).

**Figure 4:**
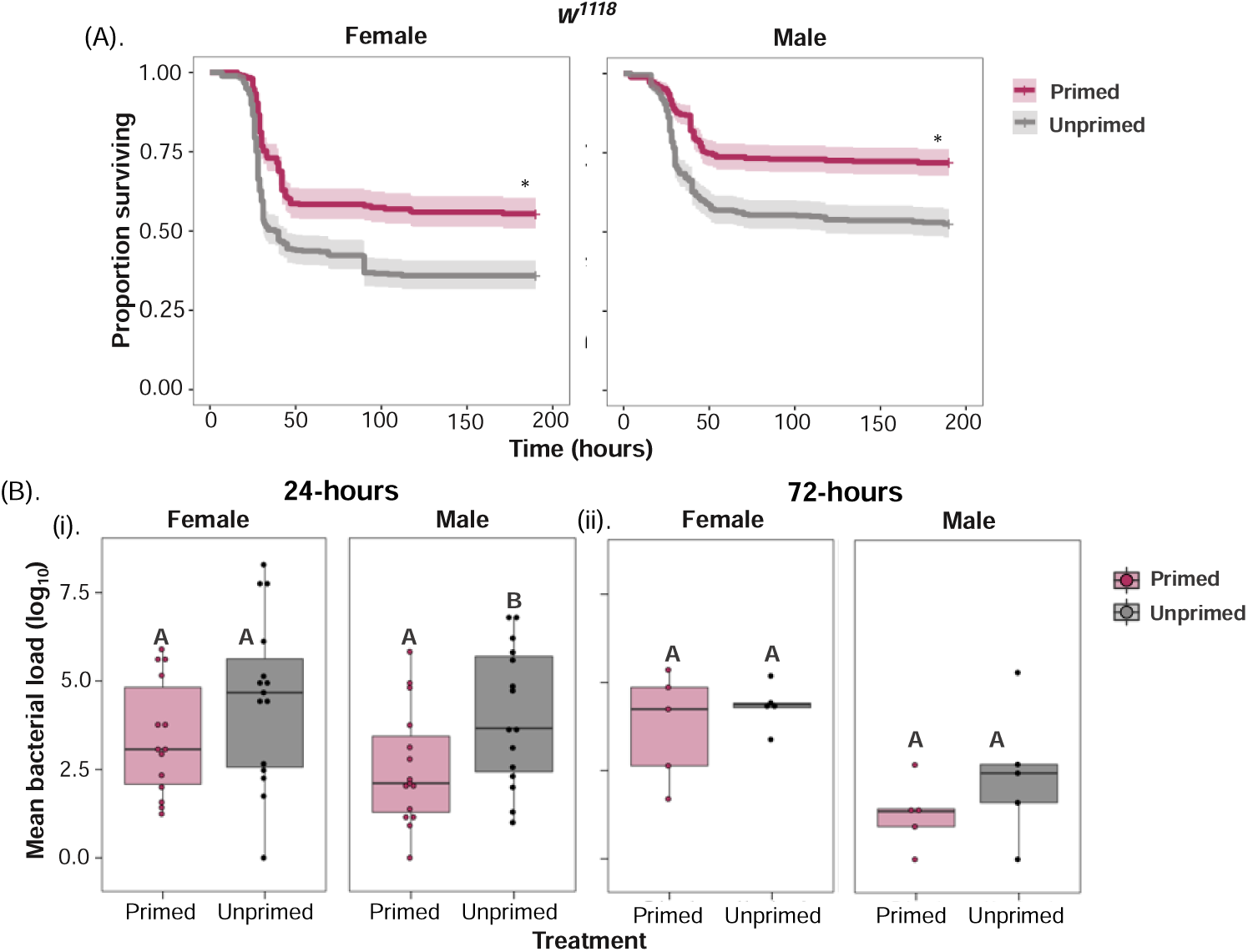
Bacterial loads of primed and unprimed *w^1118^* flies. (A). Survival curves of primed and unprimed control *w^1118^* flies (B). Internal bacterial load (n=20-22 vial with 5-7 flies in each vial/sex/treatment) after (i) 24-hours and (ii) 72-hours following *P. rettgeri* infection. Different letters in panel-B denotes primed and unprimed individuals are significantly different, tested using Tukey’s HSD pairwise comparisons, analysed separately for each timepoint and sex combination. The error bars in panel B represent standard error.

### Priming following oral infection reduces bacterial shedding and transmission in males

Apart from increasing survival during infections, immune priming may also have consequences to pathogen spread and transmission (Tate, 2016). Despite having enhanced survival, the primed individuals may also extend the infectious period or their pathogen burden may be lower, which would lead to less severe epidemics (Tate, 2016). To investigate how immune priming affects epidemiological parameters we tested whether primed individuals have reduced bacterial shedding and spread. Given the oral-faecal nature of bacterial transmission and that our previous results all related to systemic infections, we first established that a survival benefit fo priming also occurs under oral infection. Following an initial oral exposure to a heat-killed *P. rettgeri* culture, after a 72-hour period we exposed female and male *w^1118^*flies orally to a lethal dose of live *P. rettgeri*. Primed *w^1118^* males, but not females, showed increased survival decreased internal bacterial loads after priming via the oral route of infection. (**Fig.** 5A and 5B, **Table**-S7 and S8).

**Figure 5:**
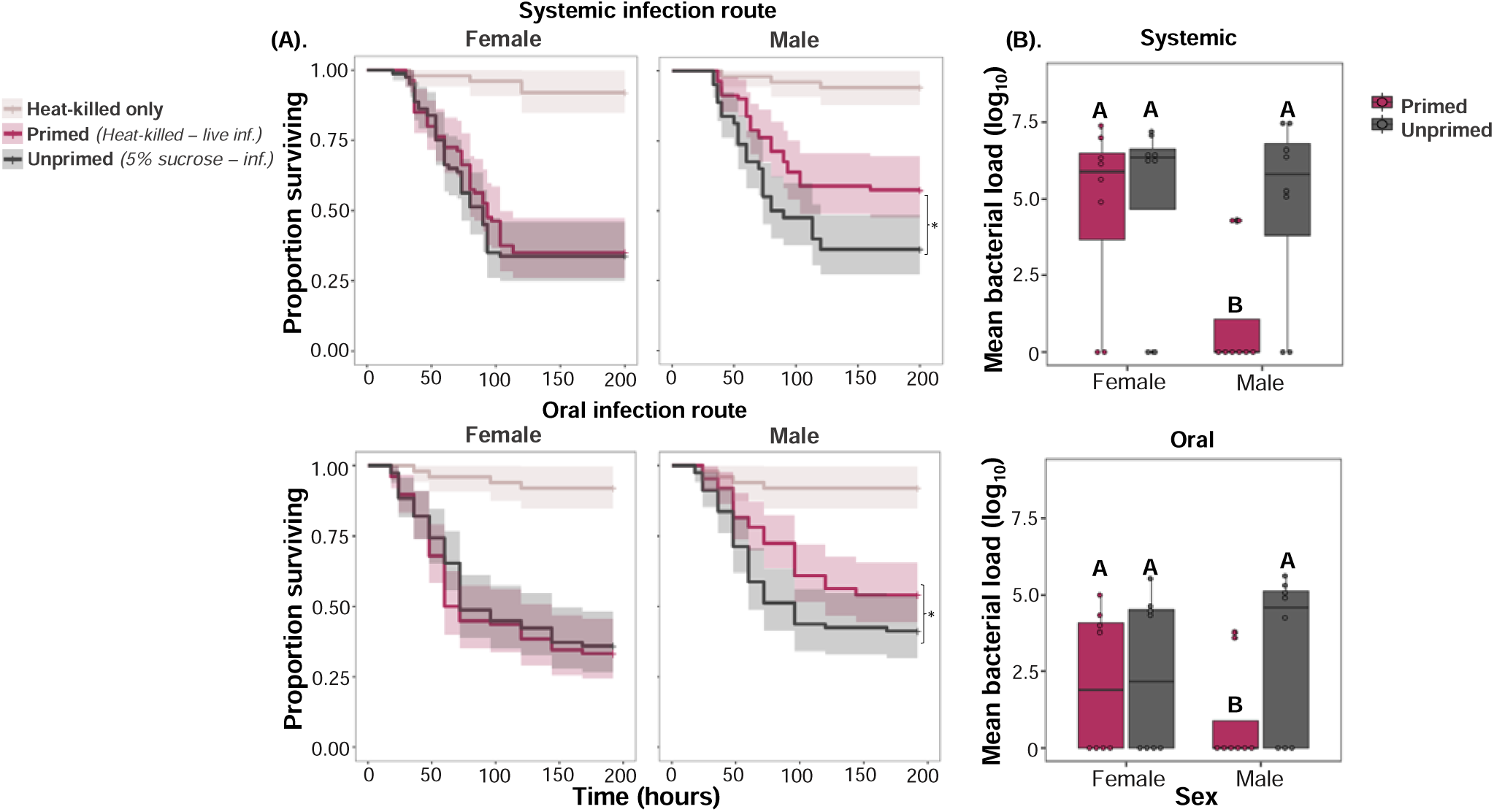
Priming occurs in males but not females during oral infections. (A) Survival curves of male and female wildtype (*w^1118^*) flies with (i) systemic (ii) oral priming where flies initially exposed to heat-killed *P. rettgeri* or unprimed flies exposed to a sterile solution in the first exposure, this is followed by live *P. rettgeri* pathogen after 72-hour time gap [systemic infection dose = 0.75 OD (∼45 cells/fly) and oral *P. rettgeri* infection dose = 25 OD_600_]. (n= 7-9 vials/sex/treatment/infection route) (B). Mean bacterial load measured as colony forming units at 24 hours following (i) systemic and (ii) oral priming, followed by live *P. rettgeri* infection for male and female *w^1118^*. Different letters in panel-B denotes primed and unprimed individuals are significantly different, tested using Tukey’s HSD pairwise comparisons.

Given this priming effect on bacterial loads (Fig 5B) we hypothesised that priming could directly impact the amount of bacterial shedding, and therefore have a direct impact on pathogen transmission. We measured the shedding of single flies 4-hours after infection by live *P. rettgeri* exposure. We found that male flies previously primed with a heat-killed bacterial inoculum shed less bacteria than unprimed flies. However, both primed and unprimed females showed increased bacterial shedding following oral *P. rettgeri* infection (**Fig.** 6A, **Table**-S10). These effects on bacterial shedding are likely to have a drirect effect on the spread of infection in groups of individuals. In transmission assays, primed donor males also spread very little pathogen upon re-infection with oral *P. rettgeri*, compared to the unprimed male treatment where successful transmission was detected recipient flies (**Fig.** 6B, **Table-**S10). However, both primed and unprimed females showed equivalent levels of bacterial spread following oral *P. rettgeri* infection (**Fig.** 6B, **Table-**S10).

**Figure 6:**
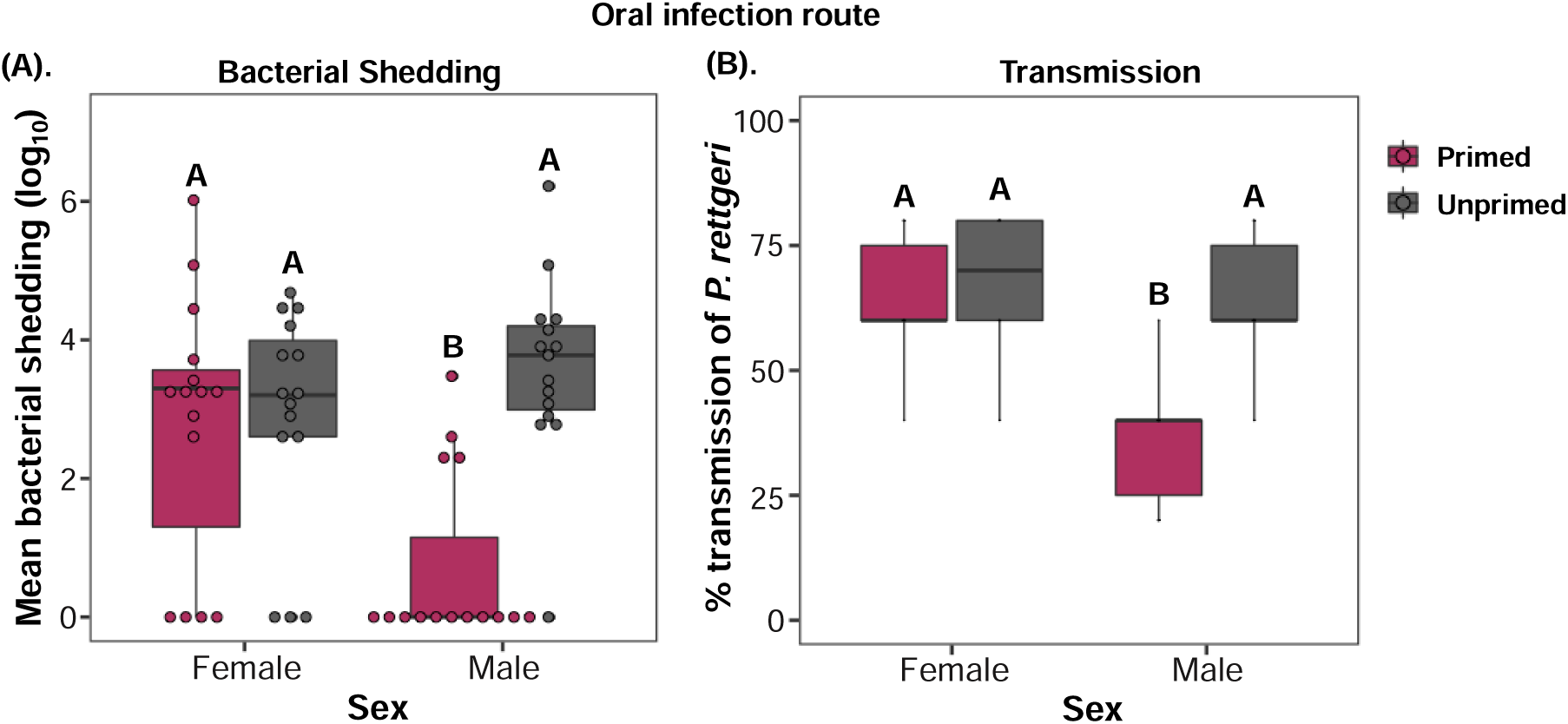
The effects of priming on bacterial shedding and transmission. (A). Bacterial shedding after 4-hours following oral priming (initial heat-killed exposure) and infection with live *P. rettgeri* (OD_600_=25) for males and females (n=15 individual flies per treatment). (B) percentage transmission of measurable bacterial loads for male and female *w^1118^* flies (recipient-flies) over the first 4-hours following exposure with infected donor flies (n=6 independent transmission assays). Different letters denote primed and unprimed individuals are significantly different, tested using Tukey’s HSD pairwise comparisons.

While multiple traits, including host activity levels and contact rates may influence pathogen transmission (Siva-Jothy and Vale, 2021; White et al., 2020), we did not find any difference in the locomotor activity of primed and unprimed flies (Fig 7). Our results therefore provide evidence that, in male flies, immune priming reduces pathogen transmission by directly decreasing bacterial shedding from infectious flies.

**Figure 7:**
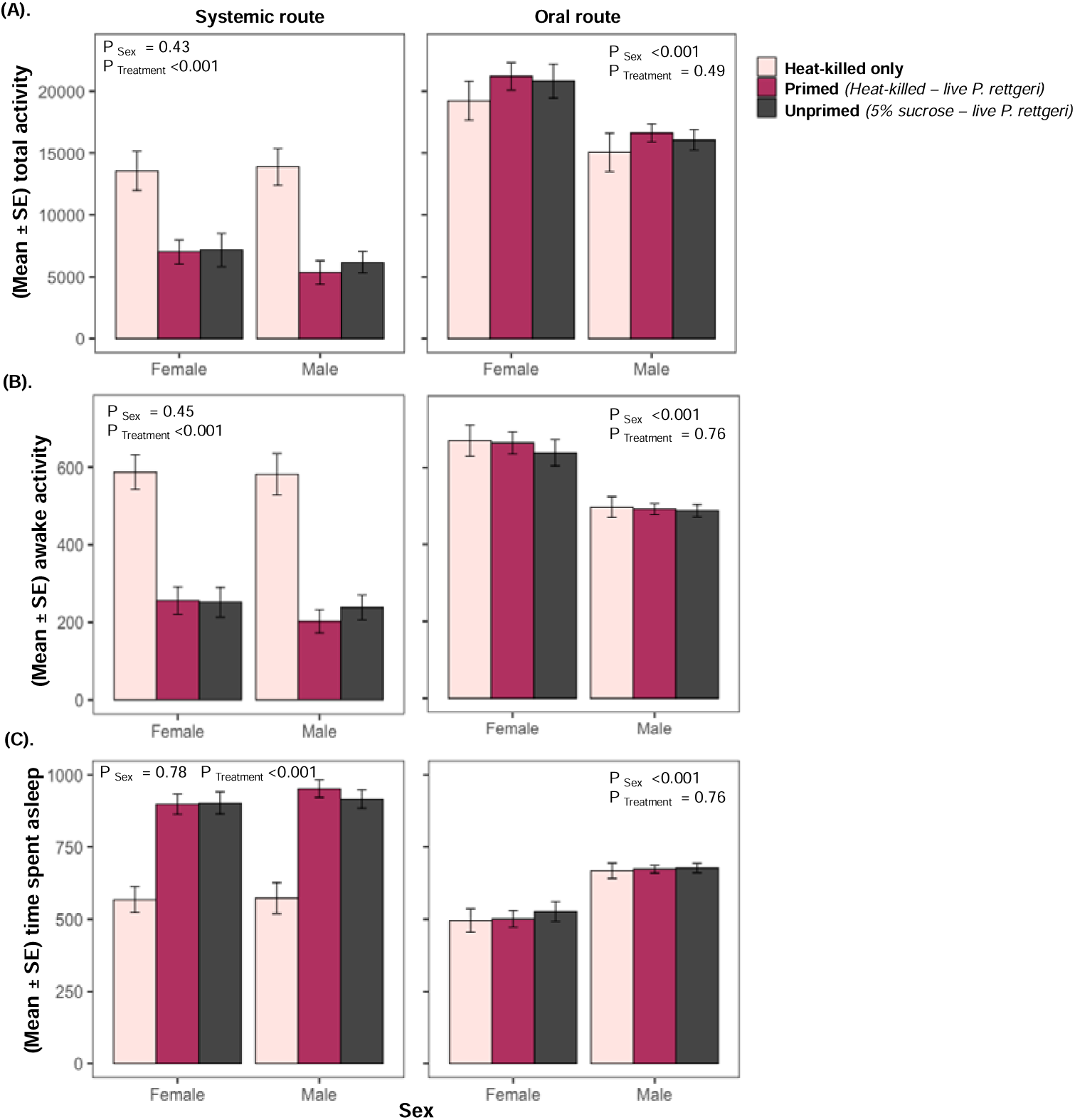
The effect of priming on fly locomotor activity. Mean ±SE total locomotor activity for males and females (n= 52 individual flies per treatment), during first 72-hours following systemic and oral priming and infection. (A) average total locomotor activity (B) average awake activity and (C) proportion of flies spent sleeping.

### The *IMD*-pathway, but not the Toll pathway, is required for immune priming during *P. rettgeri* infections

In fruit flies, the production of AMPs during antibacterial immunity is mediated by the Immune deficiency (IMD) and Toll pathways. In both pathways, pathogens are recognised by peptidoglycan receptors (PGRPs), initiating a signalling cascade that culminates in the activation of the NF-κB-like transcription factors (*Dorsal* in Toll or *Relish* in IMD), resulting in the upregulation of AMP genes. The Toll-pathway generally recognises LYS-type peptidoglycan found in gram-positive bacteria and fungi. The IMD-pathway recognises DAP-type peptidoglycan found in gram-negative bacteria and produces AMPs such as *Diptericins, Attacins* and *Drosocin* among others. AMPs work with a high degree of specificity, so that only a small subset of the total AMP repertoire provides the most effective protection against specific pathogens (Hanson et al., 2019),although this specificity has been shown to be greatly reduced during aging (Shit et al., 2022).

The inducible AMPs regulated by the IMD signalling pathway play a crucial role in resisting gram-negative bacterial infection such as *P. rettgeri* (Lemaitre and Hoffmann, 2007). Therefore, we wanted to investigate whether the IMD signalling pathway and IMD-responsive AMPs contribute to immune priming. To address this, we used several transgenic fly lines (CRISPR knockouts and UAS-RNAi knockdowns) with functional absence of or knockdown of different regulatory and effector components of the IMD-signalling pathway. First, we used a *Relish* loss-of-function mutant *Rel^E20^*, a key regulator of the IMD immune response. As expected, we found that *Relish* mutants did not show any bacterial clearance, died at faster rate, and therefore did not show any benefit of a priming treatment (**Fig**. 8A-ii for survival and **Fig**. 8B-ii for bacterial load, **Table**-S11 for survival and **Table**-S12 for bacterial load). Loss-of—function of *spätzle **(spz),*** a key regulator in the *Toll* pathway, showed enhanced survival following initial heat-killed exposure, and their mortality rates were the same as in controls (*w^1118^*) (**Fig**. 8A-iii, **Table**-S11) indicating that Toll-pathway does not contribute to priming during *P. rettgeri* infection (**Fig**. 6A-iii, **Table**-S11).

**Figure 8.**
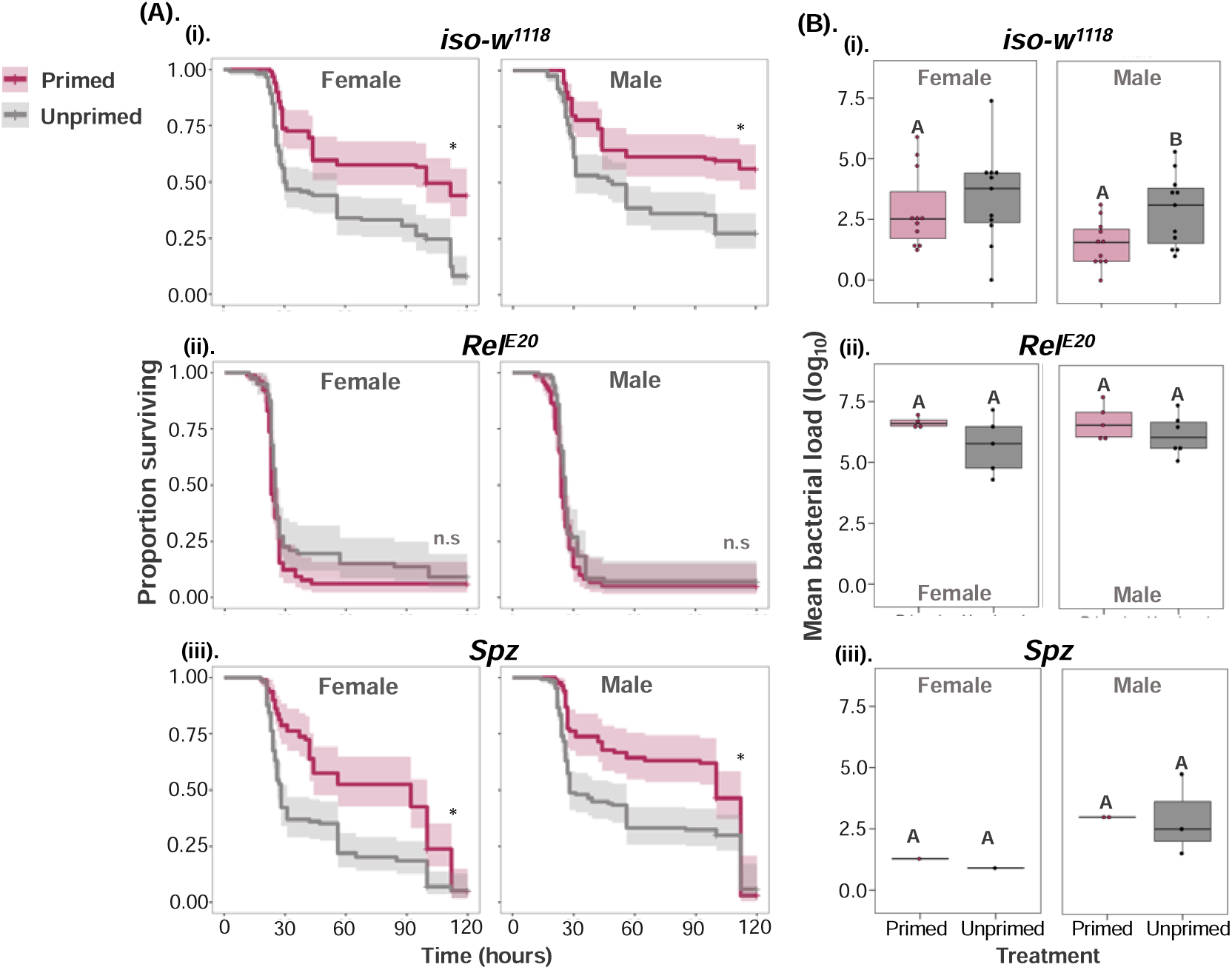
Priming requires the IMD but not the Toll pathway. (A): Survival curves for control flies and flies lacking different innate immune pathway components in females and males (i) control *iso*-*w^1118^* (ii) *Rel^E20^*-IMD-pathway transcription factor (iii) *Spz*-Toll pathway regulator in both males and females (B) (i-iii) bacterial load measured after 24-hours post-secondary pathogenic exposure (n=7-9 vial/sex/treatment/transgenic lines). Different letters in panel-B indicate that primed and unprimed individuals are significantly different, tested using Tukey’s HSD pairwise comparisons, analysed separately for each timepoint and sex combination. The error bars in panel B represent standard error.

### Expression of IMD-regulated *Diptericin B* in the fat body is required for priming

Given the important role of the IMD pathway for the priming response (Fig. 5A), next, we tested whether mutants with defective IMD signalling or unable to produce specific antimicrobial peptides were capable of immune priming. We first used a ΔAMP transgenic line, which lacks most of the known *Drosophila* AMPs (10 AMPs in total). ΔAMPs flies are extremely susceptible to the majority of microbial pathogens, including gram-negative bacteria (Hanson et al., 2019), and we confirmed that both primed and unprimed ΔAMP flies succumb to death at a similar rate, and both primed and unprimed ΔAMP flies also exhibited elevated bacterial loads (measured 24-hours after the secondary pathogenic exposure) (**Fig**. 9A-i for survival, **Fig**. 9B-i for bacterial load, **Table**-S11 and S12, relative to control w^1118^ flies – compare with previous figure Fig. 8B-i), showing that AMPs are needed for the priming response against *P. rettgeri*. To investigate which AMPs are required for priming against *P. rettgeri* infection, we used a Group-B transgenic line, lacking major IMD regulated AMPs (including *Diptericins* and *Attacins* and *Drosocin*) but have all upstream IMD signalling intact. Primed Group*-*B flies showed mortality similar to unprimed Group-B flies (**Fig**. 9A-ii, **Table** S11) and exhibited increased bacterial loads across both the sexes irrespective of being primed or not (**Fig**. 9B-ii, **Table** S12). Thus, despite being able to produce other AMPs, removal of IMD-regulated AMPs completely eliminated the priming effect, indicating that AMPs regulated by the IMD-pathway are required for immune priming against *P. rettgeri* infection.

**Figure 9:**
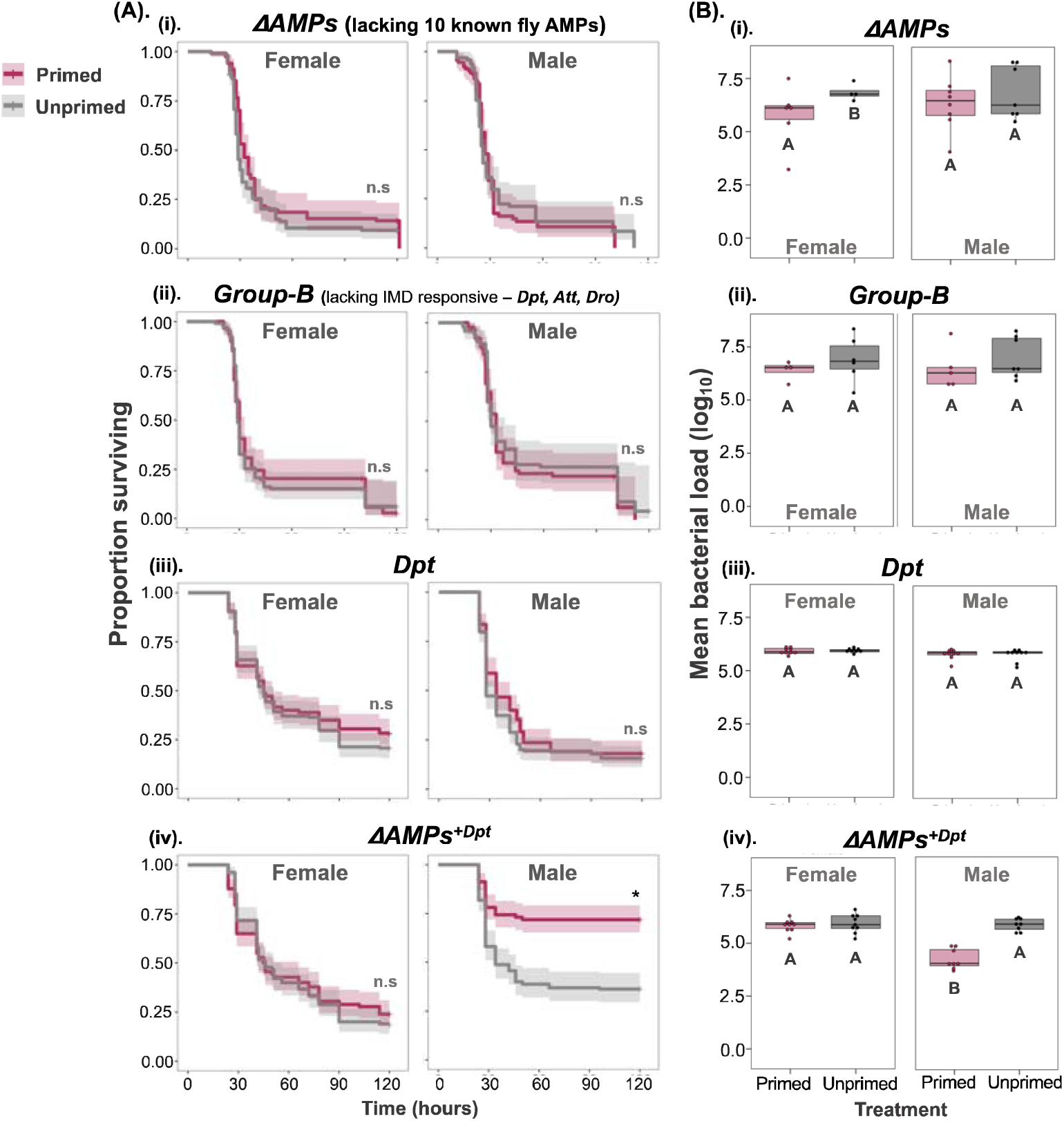
Priming requires specific AMP expression. (A). Survival curves for CRISPR/Cas9 AMP mutants (i) Δ*AMP* fly line: lacking all known 10 fly AMPs – (ii) *Group-B:* lacking major IMD pathway AMPs (iii) *Dpt* knockout and (iv) Δ*AMP^+Dpt^*: lacking all AMPs except *Dpt*. (B) internal bacterial load quantified after 24-hours post-secondary exposure (n=7-9 vials with 10-15 flies in each vial/sex/treatment/transgenic lines). Different letters in panel-B denotes primed and unprimed individuals are significantly different, tested using Tukey’s HSD pairwise comparisons, analysed separately for for each fly line and sex combination. The error bars in panel B represent standard error.

Since *Diptericins* have been shown previously to play a key role in defense against *P. rettgeri*, we then used a *Dpt* mutant (lacking *Diptericin*-*A* and *B*) and *AMPs^+Dpt^* transgenic fly lines (flies lacking all known AMPs except *Diptericin*) to test whether *Diptericins* are required and sufficient for priming in both females and males. The survival benefit of priming disappeared in flies lacking *Dpt* across both females and males and both primed and unprimed *Dpt* mutants exhibited increased bacterial load (**Fig**. 9B-iii, **Table** S12). However, the priming response was recovered completely in male flies that lacked other AMPs but possessed functional *Diptericins* (*AMPs^+Dpt^*). However, the same effect was not seen in females (**Fig**. 9A-iv for survival, **Fig**. 9B-iv for bacterial load; **Table** S11 for survival, **Table** S12 for bacterial load).

In response to gram-positive *Streptococcus pneumoniae* infection, previous work described the role of haemocytes in immune priming through increased phagocytosis (Pham et al., 2007). Subsequent work has also shown that reactive oxygen species (ROS) burst from haemocytes is important for immune priming during *Enterococcus faecalis* infection (Chakrabarti and Visweswariah, 2020). Since our results pointed to a *Diptericins* being required for immune priming against *P. rettgeri*, we wanted to determine if *Diptericin* expression in either the fat body or haemocytes was more important for immune priming. Using-tissue specific *Diptericin-B* UAS-RNAi knockdown, we found that male flies with knocked-down *DptB* in fat bodies no longer showed immune priming compared to the background or control iso-*w^1118^*, while knocking down *DptB* in haemocytes resulted in a smaller but still significant increase in survival following an initial exposure (**Fig**. 10, **Table**-S13). This would therefore support that immune priming requires *DptB* expression in the fat body, while haemocyte-derived *DptB* is not important for priming. It is worth noting that it is possible that haemocytes contribute to immune priming through phagocytosis or melanisation, or via cross-talk with the IMD pathway, and this remains a question for future research.

**Figure 10:**
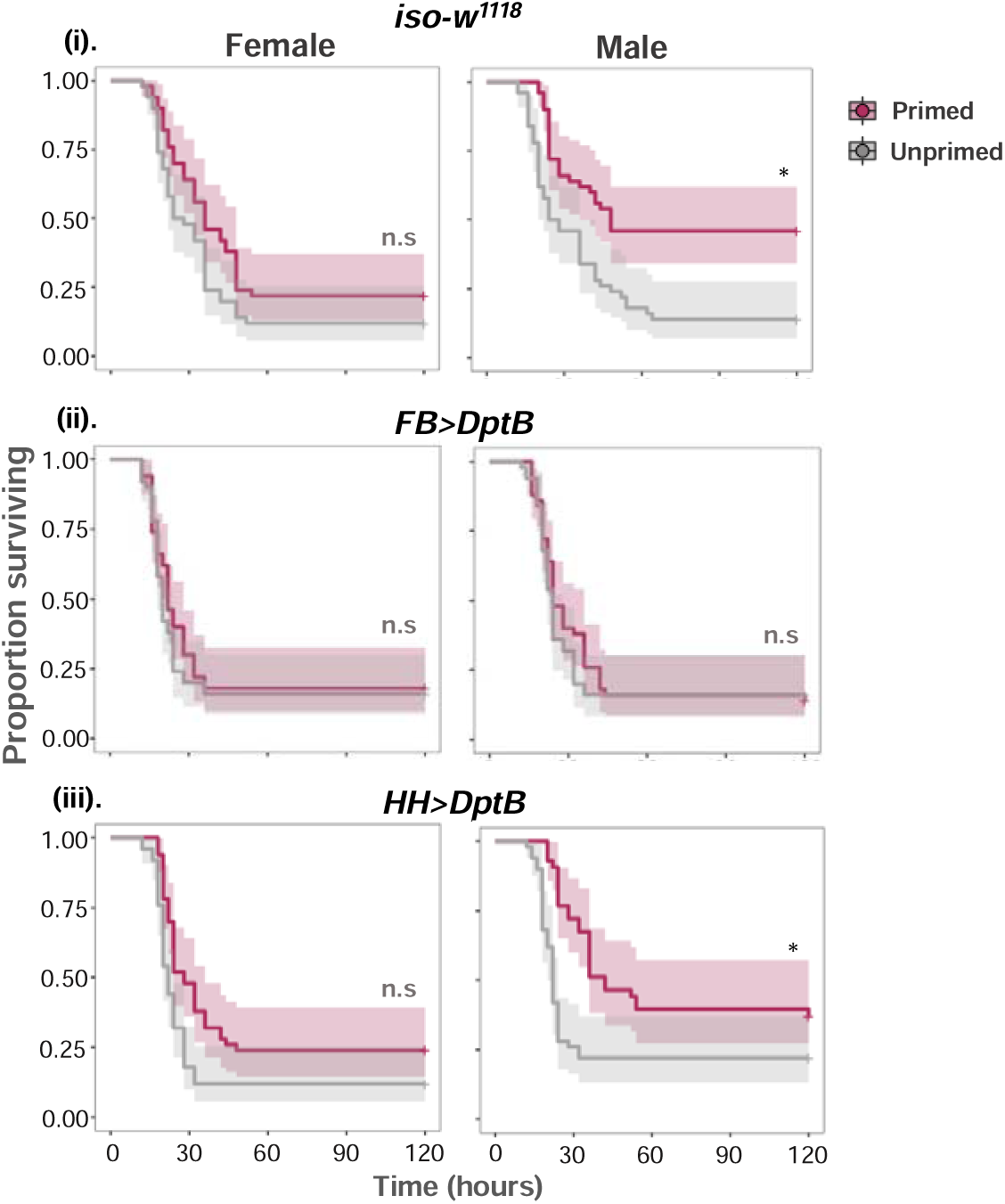
Survival curves tissue-specific Dpt knockdowns. (i) control *iso-w^1118^* and *UAS-RNAi* mutants - (ii) *FB>DptB* RNAi knock down of *DptB* in fat body and (iii) *FB>DptB* RNAi knock down of *DptB* in haemocytes. (n=∼50 flies/sex/treatment/fly lines).

### Priming is not an outcome of constant *Dpt* upregulation

As our results indicated that AMPs, especially *Diptericins* play a key role in *Drosophila* priming against *P. rettgeri*, we wanted to test if the priming effect we had observed was a result of constant upregulation of AMPs after initial heat-killed exposure allowing rapid bacterial clearance during the secondary exposure, or if AMP expression returned to a baseline level within the 96-hours between priming and the live infection. To do this, we measured *Dpt* gene expression at 18-hours and 72-hours following initial heat-killed exposure, and then 12-hours, 24-hours and 72-hours following secondary live *P. rettgeri* infection. *DptB* expression increased 18-hours after initial heat-killed exposure but returned to baseline levels by 72-hours (**Fig.** 11A and 11B, **Table**-S14) indicating that flies did mount an immune response to the heat killed bacteria, but that this was resolved by the time they were infected with the live bacteria at 96-hours.

**Figure 11:**
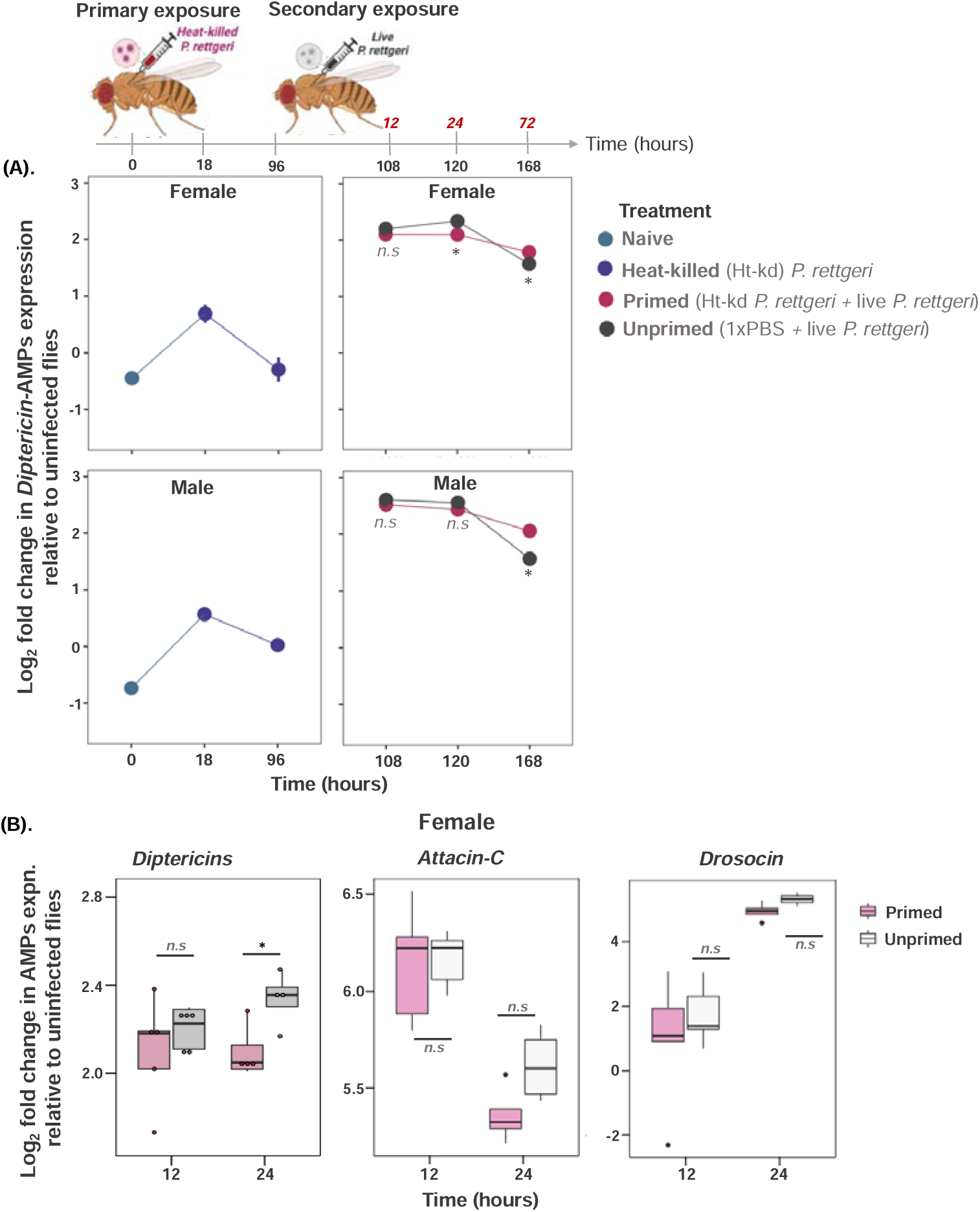
Antimicrobial gene expression following priming. (A) Mean±SE (standard error) of *Diptericin*-AMPs expression at different time points (hours) for females and males wild-type w^1118^ flies with treated with - naïve or unhandled flies; flies exposed to initial heat-killed *P. rettgeri*; primed flies that are initially exposed to heat-killed P. rettgeri followed by challenge with *P. rettgeri*; unprimed flies that receive 1xPBS during primary exposure followed by live *P. rettgeri* during secondary exposure [n=15-21 flies (3 flies pooled) /sex/treatment/timepoint]. (B). *Diptericin (A+B), Attacin-C* and *Drosocin* AMPs expression 12-hours and 24-hours after exposure to live *P. rettgeri* in w^1118^ female flies. Asterisks ‘*’ indicates that primed and unprimed individuals are significantly different (p<0.05). The error bars represent standard error.

Further, 72-hours following a lethal secondary live *P. rettgeri* infection, we found that primed *w^1118^* females and males showed increased *DptB* levels compared to unprimed individuals (**Fig.** 12A and 12B). The increased *DptB* expression also resulted in increased bacterial clearance in males (see **Fig**. 4Bi and 8Bi). In case of females, despite higher *Dpt* levels at 72-hours following secondary live infection in primed flies, we did not detect any difference in bacterial clearance between primed and unprimed flies (see **Fig**. 4Bi and 8Bi). We also tested the expression pattern of other IMD-responsive AMPs such as *Attacin-C* and *Drosocin* in females. Overall, we found that the expression of *Attacin*-*C* and *Drosocin* was not different between primed and unprimed females (**Fig.** 11C for *AttC and Dro*, **Table**-S14).

**Figure 12:**
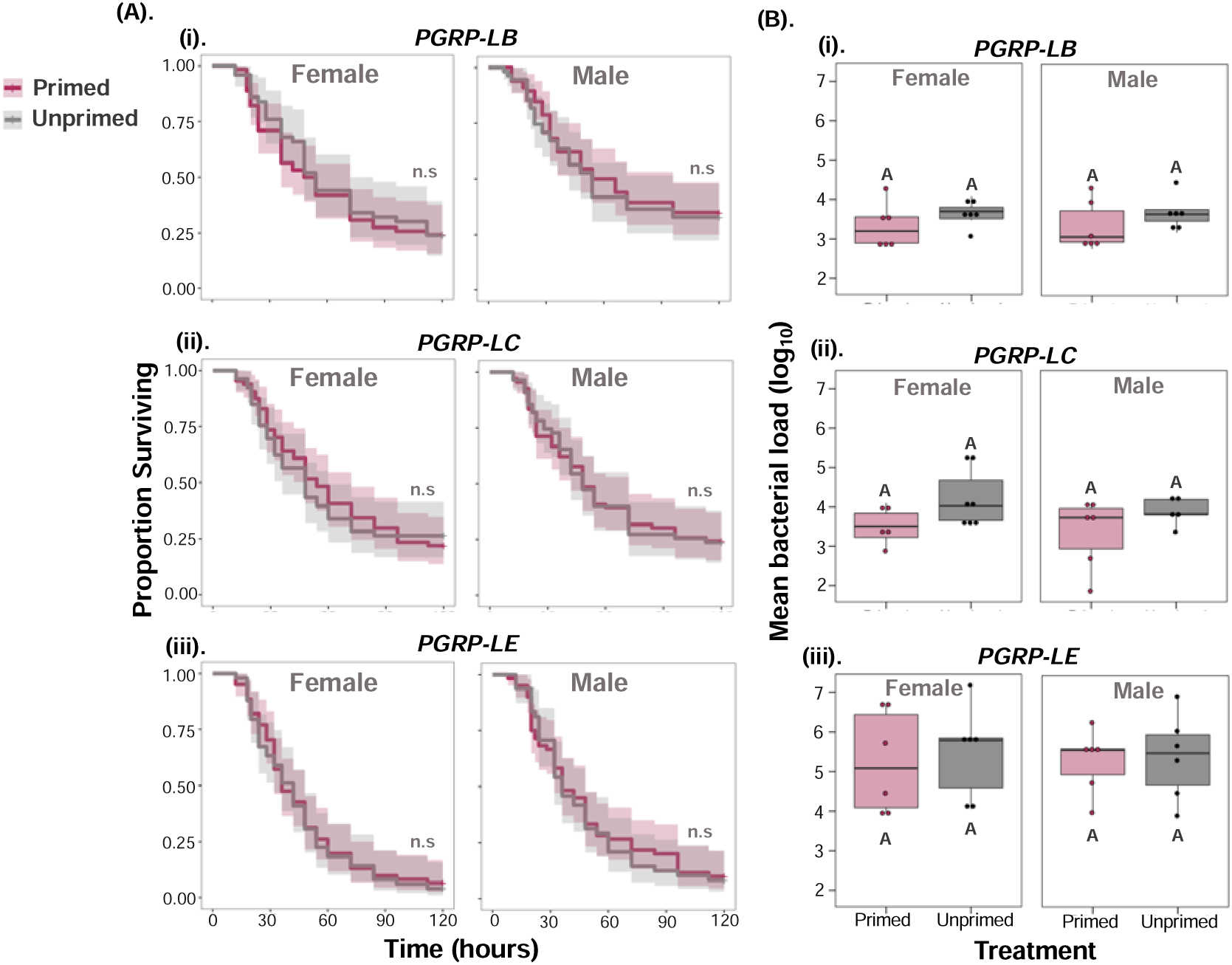
(A). Survival curves for males and females of Loss-of-function in *PGRPs* (peptidoglycan recognition proteins) of IMD **(i)** *PGRP-LB* **(ii)** *PGRP-LC* and **(iii)** *PGRP-LE* **(B).** internal bacterial load quantified after 24-hours post-secondary exposure in female and male flies (n=7-9 vials with 10-15 flies in each vial/sex/treatment/ fly lines). Different letters in panel-B denotes primed and unprimed individuals are significantly different, tested using Tukey’s HSD pairwise comparisons for each fly line and sex combination. The error bars represent standard error.

### Regulation of IMD by PGRPs is required for immune priming

Following gram-negative bacterial infection, DAP-type peptidoglycans from gram-negative bacteria are recognised by the peptidoglycan receptors *PGRP-LC* (a transmembrane receptor) and *PGRP-LE* (a secreted isoform and an intracellular isoform), which then activate the IMD intracellular signalling cascade (Lemaitre and Hoffmann, 2007; Valanne et al., 2022). Immune regulation is achieved by several negative regulators, including *PGRP-LB*, a secreted peptidoglycan with amidase activity, that break down peptidoglycans into smaller, less immunogenetic fragments(Lemaitre and Hoffmann, 2007; Valanne et al., 2022). *PGRP-LB* has also been shown to be important for transgenerational immune priming in *Drosophila* against parasitoid wasp infection (Bozler et al., 2020). Given the role of DptB in the observed priming benefit, we therefore decided to investigate the role of *PGRPs* in the immune priming we observed during *P. rettgeri* infection.

To address this, we used fly lines with loss-of-function in *PGRP-LB*, *LC* and *LE*. Regardless of which *PGRP* was disrupted, we observed that flies were no longer able to increase their survival following an initial exposure (**Fig**. 12A, **Table**-S15). However, unlike similar outcomes with *Relish* or Δ*AMP*, here the lack of priming was not driven by an inability to clear bacterial loads, as microbe loads in *PGRP* mutants were ∼100-fold lower than in *Relish* or AMP mutants (**Fig**. 12B, **Table**-S16; also compare **Figs**. 8B-ii and 9B-i. Further, Dpt expression compared to the *w^1118^*control was either similar (PGRP-LC, PGRP-LE) or higher (PGRP-LB), as expected, in both primed and unprimed flies (**Fig.** 13, **Table**-S17). This suggests that *Diptericin* expression is required, but not sufficient, for successful priming, which requires adequate regulation by *PGRPs*. Which specific molecular signal modifies *PGRP-mediated* regulation of *Diptericin* expression following an initial priming challenge is unclear, but must lie upstream of the IMD pathway and the fat body.

**Figure 13:**
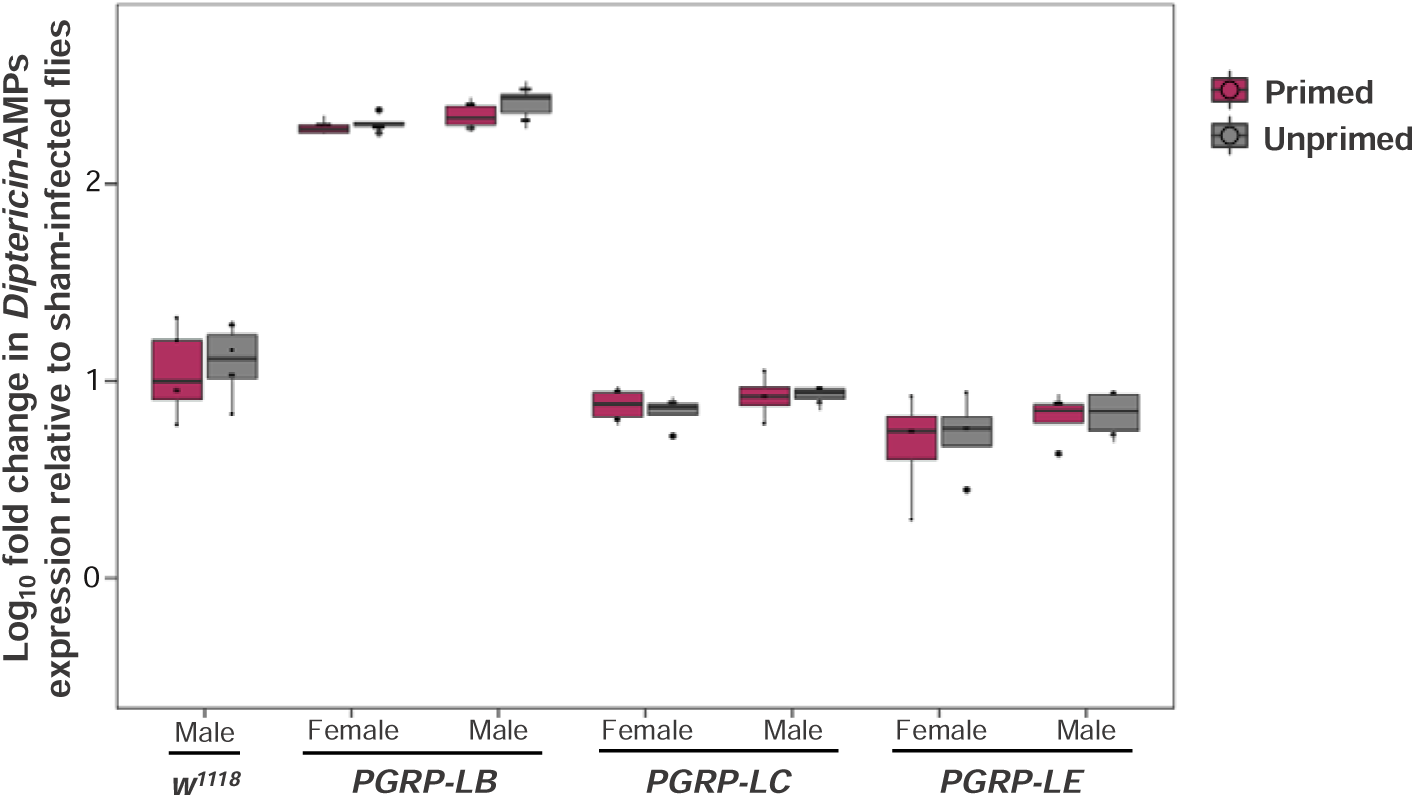
*Diptericin-*AMPs expression 24-hours after exposure to live *P. rettgeri* in male and female control *w^1118^* flies and flies with loss-of-function in different *PGRPs*, *PGRP-LB, PGRP-LC & -LE*.

## Discussion

The observation of immune priming in invertebrates underlines the importance of whole organism research in immunity in lieu of a purely mechanistic approach to immunology, highlighting that the same phenomenology can originate in very different mechanisms (Arch et al., 2022; Little et al., 2005). There is now substantial evidence of diverse priming responses occurring within the same invertebrate life stage (larvae or adults), across life stages (from larvae to adults), or even between generations (parents to offspring) (Arch et al., 2022; Contreras-Garduño et al., 2016; Khan et al., 2016; Sheehan et al., 2020). The survival benefit of priming has been observed in a range of arthropod taxa, including Dipterans: fruit flies (Pham et al., 2007), mosquitoes (Ramirez et al., 2015); Coleopterans: flour beetles (Khan et al., 2016; Moret and Siva-Jothy, 2003); Lepidopterans: the greater wax-moth (Fallon et al., 2011); Hymenopterans: bumblebee (Sadd and Schmid-Hempel, 2006); Crustaceans: water fleas (Little et al., 2003) and Arachnids: spiders and scorpions (Gálvez et al., 2020). Thus, substantial experimental evidence in both lab adapted and wild-caught arthropods suggests that immune priming is a widespread phenomenon, and is predicted to have a profound impact on the outcome of host–pathogen interactions, including infection severity and pathogen transmission (Tate and Rudolf, 2012; Tidbury et al., 2012; Tate, 2016; Gomes et al., 2022).

In the present work, we investigated the occurrence, generality, specificity and mechanistic basis of immune priming in fruit flies when infected with the gram-negative pathogen *Providencia rettgeri*. We present evidence that priming with an initial non-lethal bacterial inoculum results increased survival after a secondary lethal challenge with the same live bacterial pathogen. This protective response may last at least two weeks after the initial exposure, is particularly strong in male flies, and occurs in several genetic backgrounds. We show that the increased survival of primed individuals coincides with a transient decrease in bacterial loads, and that this is likely driven by the expression of the IMD-responsive AMP *Diptericin-B* in the fat body. Further, we show that while *Diptericin* is required as the effector of bacterial clearance, it is not sufficient for immune priming, which requires the regulation by at least three PGRPs (*PGRP-LB, PGRP-LC*, and *PGRP-LE*). Therefore, despite having an intact IMD signalling cascade, and being able to express *Diptericins*, flies lacking any one of *PGRP-LB, LC* or *LE* were not capable of increasing survival following an initial sublethal challenge. Together, our data indicates that PGRPs are necessary for regulating immune priming against *P. rettgeri*.

These results are largely consistent with other work showing that the pathways that signal pathogen clearance may not be the same that underlie the signalling of the priming response. For example, Cabrera et al investigated priming with the Gram-positive *E. faecalis* and found that while the Toll pathway was required for responding to a single infection, Toll signalling was dispensable for immune priming (Cabrera et al., 2023). By contrast, IMD-deficient flies were still able to clear single *E. faecalis* infections (due a functional Toll response) but were no longer able to increase survival following an initial exposure. That is, in support of our results here, the IMD played a distinct role in signalling priming that was independent of its role in clearance. Further, previous work found that the Toll pathway is required (though not sufficient) for priming in response to gram-positive bacterial *Streptococcus pneumoniae* infection (Pham et al., 2007) and here we find that the Toll pathway is not required for a functional immune priming response.

In response to infection, the expression of several AMP genes in *Drosophila* increases and then drops once the infection threat is resolved or controlled. Previous studies have shown that this occurs within a period of few hours during bacterial infections (Lemaitre et al., 1997), and that in many cases, this increased AMP expression underlies the immune priming response (Arch et al., 2022; Wen et al., 2019). In other host-pathogen systems, genome wide insect transcriptome studies have identified upregulation of several AMPs in primed individuals, for example, *Attacins, Defensins and Coleoptericins* in flour beetles (Ferro et al., 2019; Greenwood et al., 2017), *Cecropin, Attacin, Gloverin, Moricin* and *Lysozyme* in silkworms (Yi et al., 2019), *Gallerimycin and Galiomicin* in wax-moths (Bergin et al., 2006) and finally, *Cecropin* in tobacco moths and fruit flies (Roesel et al., 2020; Chakrabarti and Visweswariah, 2020). However, there are also examples where innate immune priming involves the downregulation of AMP expression, such as priming with the gram-positive *E. faecalis* (Cabrera et al., 2023).

One key aspect of our results is that immune priming was not the result of continued upregulation of *Diptericin* following the initial exposure to heat-killed bacteria. Instead, we observed that the initial immune response to heat-killed bacteria had already been resolved at 72-hours, before the secondary exposure to a lethal infection at 96-hours (see **Fig**. 11A and B). This second up-regulation of AMP expression was at least 10-times higher than the response to heat-killed bacteria, and was initially similar between both primed and unprimed flies (exposed to a sterile solution in the first exposure). It was only at 72-hours following the second lethal exposure that we observed significantly higher expression of *Diptericin* in primed flies. This difference appears to arise because unprimed flies show a faster resolution of the immune response compared to primed flies; that is, the expression of *Diptericin* shows a faster decline between 24-hours and 72-hours post-exposure in unprimed compared to primed flies (see **Fig**. 11A and B). These patterns of gene expression provide a partial explanation for the increased survival following priming, but they do not explain why primed individuals had lower bacterial loads at 24-hours after exposure to the second lethal infection, as at this timepoint we did not detect any differences in *Diptericin* expression between primed and unprimed flies. It is also unclear why primed females showed increased expression of *Diptericin* after 24-hours, but do not show any reduction in bacterial loads at this timepoint following priming, in the way male flies did.

The sex differences we observed in priming reflect a larger pattern of sexual dimorphism in immunity present in most including *Drosophila* (Belmonte et al., 2020; Klein and Flanagan, 2016; Simon, 2005). Here we found that males showed better bacterial clearance after initial exposure to heat-killed *P. rettgeri* which enabled them to experience enhanced survival compared to females, who exhibited higher bacterial loads and greater mortality. One possibility for the observed sex-differences could relate to different nutritional demands and metabolic activities in female *Drosophila*. For instance, female fruit flies are often able to reallocate resources in accordance with their reproductive demands, as observed in terminal investment during infection (Camus et al., 2019; Martínez et al., 2020; Hudson et al., 2020). Studies from other insects suggest that inducing priming responses can directly reduce the reproductive fitness in mosquitoes (Contreras-Garduño et al., 2014), wax moth (Trauer and Hilker, 2013), and mealworm beetles (Zanchi et al., 2011). There are therefore potential trade-offs between investment in reproductive effort and investment in stronger immune responses following priming. Given that all females used in this study were mated, it is possible that such trade-offs forced a reallocation of resources towards reproduction, thereby reducing the observed magnitude of the priming response in females. Future studies may consider comparing priming responses in females with different reproductive states in order to test whether immune priming is costlier for female *Drosophila*.

A subsidiary finding of this work was the effect of the endosymbiont *Wolbachia* on immune priming. How endosymbionts that are widespread among insects are likely to influence innate immune priming is a topic of considerable interest (Prigot-Maurice et al., 2022). There is abundant evidence that *Drosophila* carrying the endosymbiont *Wolbachia* are better able to survive infections, especially viral infections (Hedges et al., 2008; Teixeira et al., 2008; Martinez et al., 2014; Vanika Gupta et al., 2017). In this case, we observed that the priming response that was present in male flies cleared of *Wolbachia* disappeared in males carrying the endosymbiont. This effect is unlikely due to a direct effect of *Wolbachia* on the ability to clear *P. rettgeri* in primed flies, as previous work has found that *Wolbachia* had no effect on the ability to suppress *P. rettgeri* during systemic infection (Rottschaefer and Lazzaro, 2012). Indeed, if priming is the result of upregulation of AMP expression, this may suggest that *Wolbachia* may be actively suppressing the expression of AMPs, thereby reducing the beneficial effects of priming. However, this hypothesis would contradict work showing that some *Wolbachia* strains upregulate the host’s immune response and result in a reduction of pathogen growth (Rancès et al., 2012; Chrostek et al., 2013). Further rigorous experimental work is therefore required to fully understand this effect of *Wolbachia* on immune priming.

Finally, it is important to consider the implications of immune priming for disease ecology and epidemiology, and particularly how it may affect pathogen transmission. If priming acts by improving bacterial clearance via increased AMP expression, as observed in the present work, we predict that priming is likely to reduce pathogen shedding at the individual level, resulting in reduced disease transmission at the population level. Epidemiological models that have incorporated priming predict that primed individuals with enhanced survival following an initial sub-lethal pathogenic exposure are less likely to become infectious upon re-infections (Tate, 2016). It remains unclear whether immune priming reduces pathogen transmissibility, varies the infectious period, or alters infection-induced behavioural changes in the host (Tate, 2016). An additional level of complexity is that there is likely to be substantial within-population genetic variation in how priming affects each of these components of pathogen transmission. Insects offer a powerful system to investigate these effects, because they rely on an innate immune system that can induce an easily measurable priming response, but further, insect are also important vectors of many infectious diseases. Immune priming has thus emerged as a provocative idea to reduce the vectorial capacity of insect vectors, thereby reducing the transmission of vector-borne pathogens (Gomes et al., 2022). A better understanding of how immune priming contributes to host heterogeneity in disease outcomes would aid our understanding of the causes and consequences of variation in infectious disease dynamics.

## Materials and Methods

### Fly strains and maintenance

Several *D. melanogaster* strains: *w^1118^* (Vienna *Drosophila* Resource Center)*, iso-w^1118^* (Bloomington *Drosophila* Stock Center), Canton-S, Oregon-R *Wolbachia^+ve^ and Oregon-R Wolbachia^-ve^ (Vanika Gupta et al., 2017).* Transgenic flies included immune mutants *Rel^E20^* (*relish - IMD* pathway regulator (Hedengren et al., 1999)) and *spz (spätzle – Toll* pathway regulator (Levashina et al., 1998), and were a gift from the Saleh lab (Pasteur, Paris). We also used the following CRISPR/Cas9 deletion lines, originally gifted by the Lematire lab (EPFL, Lausanne) (Hanson et al., 2019): (a) ΔAMPs *-* flies lacking 10 fly AMPs, (b) Group-B - flies lacking major IMD regulated AMPs including *Attacins* (*AttC^Mi^; AttD^SK1^), Drosocin (Dro^SK4^) and Diptericins (Dpt^SK12^*), (c) *Dpt^SK12^* – flies lacking *Diptericins (DptA and DptB),* and (d) ΔAMPs*^+Dpt^* – flies lacking 10 known AMPs except *Diptericins*. All the CRISPR/Cas9 mutants were generated previously from the *iso-w^1118^* genetic background using CRISPR*/*Cas9 gene editing technology to induce null mutations in the selected genes (Hanson et al., 2019).

We also used the binary GAL4/UAS system for tissue specific silencing of target genes with the RNA interference (RNAi) method. *Diptericin-B (DptB)* (Bloomington stock# *28975*) was tissue-specifically knocked down in the fat body (*w^1118-iso^,Fb-Gal4i+(P{fat}*) and in haemocytes [*w^1118-iso^,Hmldelta-Gal4; He-Gal4* – a combination of two haemocyte GAL4 drivers, the Hml-GAL4.Δ (Sinenko and Mathey-Prevot, 2004) and He-GAL4.Z (Zettervall et al., 2004)]. All fly stocks and experimental flies were maintained at 25°C ±1°C on a 12:12 hour light: dark cycle in vials containing 7ml of standard cornmeal fly medium (Lewis, 2014). For the experiments, we controlled larval density by placing 10 females and 5 males in each vial and the females were allowed to lay eggs for 48-hours. Fourteen days later, the eclosing males and females were sorted and collected and separated into group of 25 flies in each vial. Three-day old, mated individuals were used in all experiments.

### Systemic immune priming and infection assays

*P. rettgeri* was grown at 37°C in 10ml Luria broth (Sigma Ltd) overnight to reach optical density OD_600_=0.95 (measured at 600nm in a Cell Density Meter, Fisherbrand™). The culture was centrifuged at 5000 rpm for 5 min at 4°C, and the supernatant was removed, and the final OD was adjusted to 0.1 and 0.2 by using sterile 1xPBS (Phosphate buffer saline). To obtain heat-killed bacteria the dilution was incubated at 90°C for 20-30 mins (Khan et al., 2016). To ensure all bacteria were dead in the heat-treated culture, it was plated and no growth was confirmed. To prime individuals, 3-day old adults were pricked with a 0.14-mm pin (Fine Science Tools) dipped in either heat-killed bacteria for the primed treatment or in 1xPBS solution for the unprimed treatment (exposed to a sterile solution in the first exposure) in the mesopleuron region (the area situated under the wing and to the left of the pleural suture) (Prakash et al., 2021). Following this initial priming treatment, the individuals were pricked using OD_600_=0.1 live *P. rettgeri* bacteria (resulting in approximately 70 bacterial cells/fly). To test whether male and female adult flies show priming (measured as enhanced survival) with increasing time intervals between the initial heat-killed exposure and later challenge with live *P. rettgeri*, we tested several time points between the two challenges 18-hours, 48-hours, 96-hours, 1-week and 2-weeks (See **Fig**. S1 for experimental design; n=9-13 vial treatment/sex/ fly line). We used a split vial experimental design to obtain replicate matched data for both survival and bacterial load see (Prakash et al., 2023) for details. Briefly after infection each vial containing about 25 flies (of each treatment, sex and fly line combination) were divided into 2 vials for measuring (i). survival following infection (see **Fig**. S1i; 13-17 flies/combination) and (ii). internal bacterial load (see **Fig**. S1ii).

### Bacterial load quantification

To test whether the host’s ability to supress bacterial growth varies across primed and unprimed individuals, we quantified bacterial load as colony forming units (CFUs) at 24-hours after *P. rettgeri* infection for controls and transgenic flies in both sexes. Flies were surface sterilized in groups of 3-5 flies per vial in 70% ethanol for 30s and washed twice with distilled water before homogenising flies individually using micro pestles. We immediately performed serial dilutions of the homogenate with 1xPBS and plated them on LB agar plates and cultured at 29°C overnight. The following day, we counted the CFUs manually (Siva-Jothy et al., 2018) (n=9-13 vial/sex/treatment/fly mutants).

### Gene expression quantification

The expression of *Diptericins* was quantified by qRT-PCR. In parallel with the survival experiment, we randomly selected a subset of control w^1118^ individuals (both males and females) for RNA extraction, we included 15 flies [5-7 replicates of 3 flies pooled together for each treatment (primed and unprimed) for both males and females]. We randomly removed selected flies (3 flies per vial) at different time points post exposure to *P. rettgeri* (18-hours and 72-hours post-priming, 12-hours and 24-hours post-challenge). We then homogenised pools of three in 80μl of TRIzol reagent (Invitrogen, Life Technologies). Homogenates were kept frozen at −70°C until RNA extraction. We performed mRNA extractions using the standard phenol-chloroform method and included a DNase treatment (Ambion, Life Technologies).

We confirmed the purity of eluted RNA using a Nanodrop 1000 Spectrophotometer (version 3.8.1) before going ahead with reverse transcription (RT). The cDNA was synthesized from 2μl of the eluted RNA using M-MLV reverse transcriptase (Promega) and random hexamer primers, and then diluted 1: 1 in nuclease free water. We then performed quantitative RT-PCR (qRT-PCR) on an Applied Biosystems StepOnePlus machine using Fast SYBR Green Master Mix (Invitrogen) using a 10μl reaction containing 1.5L of 1:1 diluted cDNA, 5μl of Fast SYBR Green Master Mix an 3.5μl of a primer stock containing both forward and reverse primer at 1μM suspended in nuclease free water (final reaction concentration of each primer 0.35μM). For each cDNA sample, we performed two technical replicates for each set of primers and the average threshold cycle (Ct) was used for analysis. We obtained the *AMP* primers from Sigma-Aldrich Ltd; *Dpt*_Forward: 5’ GACGCCACGAGATTGGACTG 3’, Dpt_Reverse: 5’ CCCACTTTCCAGCTCGGTTC 3’, *AttC*_Forward: TGCCCGATTGGACCTAAGC, *AttC*_Reverse: GCGTATGGGTTTTGGTCAGTTC, *Dro*_Forward: ACTGGCCATCGAGGATCACC, *Dro*_Reverse: TCTCCGCGGTATGCACACAT. We used *RpL*49 as endogenous reference gene, *RpL*49_Forward: 5’ ATGCTAAGCTGTCGCACAAATG 3’, *RpL*49_Reverse: 5’ GTTCGATCCGTAACCGATGT 3’. We optimised the annealing temperature (TC) and the efficiency (Eff) of the *Dpt* primer pair was calculated by 10-fold serial dilution of a target template (each dilution was assayed in duplicate); *Dpt*: TC= 59 °C, Eff= 102%; *AttC*: TC= 60 °C, Eff= 94%; *Dro*: TC= 61 °C, Eff= 104%.

### Oral priming and infection

For oral priming and live infection, we adjusted the final concentration to OD_600_=25 (Prakash et al., 2022; Siva-Jothy et al., 2018). We initially prepared vials for oral priming by pipetting 350-400 µl of standard agar [see (Siva-Jothy et al., 2018)] onto lid of a 7ml tubes (bijou vials) and allowed it to dry. Simultaneously, we starved the experimental flies on 12ml agar vials for 4 hours. Once the agar on the lids dried, we placed a small filter disc (Whattmann-10) in the lid and pipetted 80µl of heat-killed bacterial culture (primed treatment) or 5% sucrose solution (unprimed control treatment) directly onto the filter disc. Once the agar dried, we orally exposed flies (heat-killed only, primed and unprimed treatment) by adding approximately 10 flies per vial for 18-hours and then transferred the flies onto fresh Lewis food vials. After 3-days, we again prepared the bijou vials and once the agar dried, this time we added 80µl of live bacterial culture (OD_600_=25) and exposed flies to live *P. rettgeri* for 18-hours. We then transferred flies onto fresh food vials and observed survival after oral exposure to *P. rettgeri* every 12 hours for the following 8 days.

### Measuring locomotor activity

We measured the locomotor activity of single flies (n=52 flies for each for each sex and treatment combination) during three continuous days using a *Drosophila* Activity Monitor – DAM (v2 and v5) System (Pfeiffenberger et al., 2010), in an insect incubator maintained at 25°C ± 1°C in a 12 D: 12 L cycle. We then processed the raw activity data using the DAM System File Scan Software (Pfeiffenberger et al., 2010). We analysed fly activity data using three metrics (Anderson et al., 2022; V. Gupta et al., 2017; Siva-Jothy and Vale, 2019; Vale and Jardine, 2015): total activity, the average activity during 5-min activity bouts, and proportion of 5-min bouts with zero activity (which has been defined as sleep in *Drosophila* (Shaw et al., 2000)) (**Fig.** S2-III)

### Measuring bacterial shedding

We measured the bacterial shedding of single flies (n = 8-12 flies per treatment and sex combination) at a single time point, 4-hours following overnight oral bacterial exposure. We chose this timepoint as in other work we have found that most faecal shedding of bacterial pathogens occurs within the first 4 hours, and steadily decreases by 8 hours following overnight oral infection. Following oral priming to either heat-killed bacterial culture (primed treatment) or 5% sucrose solution (unprimed treatment), flies were exposed to an oral infection with live infection with 80µl of live *P. rettgeri* culture (OD_600_=25). Following 18 hours of oral exposure, flies were placed individually into 1.5ml Eppendorf tubes with approximately 50μl of Lewis medium in the bottom of the tube for 4-hours. After 4-hours, flies were removed from the tube and the remaining content of each tube was washed with 50μl of 1xPBS buffer by vortexing thoroughly for at least 5 secs. We then plated these samples on a LB agar plates, incubated them at 29°C and counted the colonies manually after 18-hours.

### Measuring bacterial transmission

We measured transmission in groups of flies, by collecting age-matched donor and recipient *w^1118^* flies, separately for each sex. To test the effect of priming on the ability of flies to transmit *P.rettgeri*, we orally exposed 3-day old *w^1118^* flies (donor flies) with either 5% sucrose and heat-killed bacteria (primed) or 5% sucrose only (unprimed). After 96-hours the donor flies were exposed to OD_600_=25 of *P. rettgeri* (*see oral infection section above*). The infected donors were marked by cutting the corner of a fly wing. We then placed one donor and five uninfected recipient flies in 7ml bijou vials with a small amount of Lewis food on the lid for each treatment (heat-killed bacteria only and 5% sucrose exposed without live infection) and sex-combination. After 4-hours exposure we surface-sterilised and homogenized the flies and plated the homogenate to measure the presence or absence of *P. rettgeri* infection inside each recipient fly, as an indication of successful transmission of bacteria from the donors to the recipient flies. We also set up and plated flies in groups with no infection, to confirm that our measures of *P. rettgeri* prevalence reflected successful transmission from the donor flies.

### Data analysis

All statistical analyses and graphics were carried out and produced in R (version 4.2.2) using the ggplot2, coxhz and lme4 packages (Wickham, 2016; Therneau, 2015; Bates et al., 2014), and all raw data and code is available at <u>10.5281/zenodo.7624084</u> (Prakash and Vale, 2023). We analysed the survival data following systemic *P. rettgeri* infection with a mixed effects Cox model using the R package ‘coxme’ (Therneau, 2015) for different treatment groups (that is, primed and unprimed) across both the sexes and fly lines (controls and transgenic lines). We specified the model as: survival ∼ treatment * sex * (1|vials/block), with ‘treatment’ and ‘sex’ and their interactions as fixed effects, and ‘vials’ nested within each ‘block’ as a random effect for control and transgenic lines. We used ANOVA to test the impact of each fixed effect in the ‘coxme’ model. We analysed the bacterial load, measured as log_10_ bacterial colony-forming units (CFUs) at 24-hours following *P. rettgeri* infection. As bacterial load data was non-normally distributed, we log-transformed the data and analysed using a non-parametric one-way ANOVA (Kruskal-Wallis test) to test whether the ‘treatment’ groups that is, primed and unprimed individuals significantly differed in internal bacterial load for males and females of each fly lines (control and transgenic lines).

We analysed the gene expression data by calculating the ΔΔCT value (Livak and Schmittgen, 2001). We used *RpL32* as a reference gene as it was expressing steadily in our treatment and control conditions. We calculated fold change in gene expression relative to the uninfected controls to calculate ΔΔCT and used ANOVA to test whether AMP expression differed significantly between primed and unprimed treatment for males and females. Data for bacterial shedding and transmission were non-normally distributed (tested using Shapiro-Wilks test for normality) hence we performed non-parametric Wilcoxon Kruskal-Wallis tests for bacterial shedding and transmission data.

## Acknowledgments

We thank Bruno Lemaitre’s lab for generating and generously sharing the CRISPR/Cas9 AMP transgenic lines. We thank Emily Robertshaw, Srijan Seal, Saubhik Sarkar and Pavan Thunga for laboratory assistance. We thank Sveta Chakrabarti and Ashworth fly group members for helpful discussion. Finally, we thank Angela Reid, Lucinda Rowe, James King and Alison Fulton for help with media preparation. Figures *SI-1* & *SI-2*, were prepared using Biorender platform. We acknowledge funding and support from the Branco Weiss fellowship and a Chancellor’s Fellowship to PFV; a Darwin Trust PhD studentship to AP from the School of Biological Sciences, The University of Edinburgh, United Kingdom and the Ashoka University, India. For the purpose of open access, the author has applied a Creative Commons Attribution (CC BY) license to any Author Accepted Manuscript version arising from this submission.

