## Supplementary Figures and Tables for "IMD-mediated innate immune priming increases Drosophila survival and reduces pathogen transmission"

**Supplementary Information**

### Supplementary figures

#### Figure S1. Schematic representation of different priming experiments

aimed at **(I).** testing whether the length of the period between primary heat-killed exposure and the secondary pathogenic challenge affects the extent of priming **(II).** how different lab-adapted control/genetic background flies vary in priming **(III).** dissecting the role of innate immune pathways (IMD and Toll) and inducible AMPs in immune priming and **(IV).** deciphering mechanisms that bring about immune priming in Drosophila using tissue-specific fat body and haemocytes UAS^RNAi^ mutants. The experimental design for priming assays includes survival and internal bacterial load quantification.

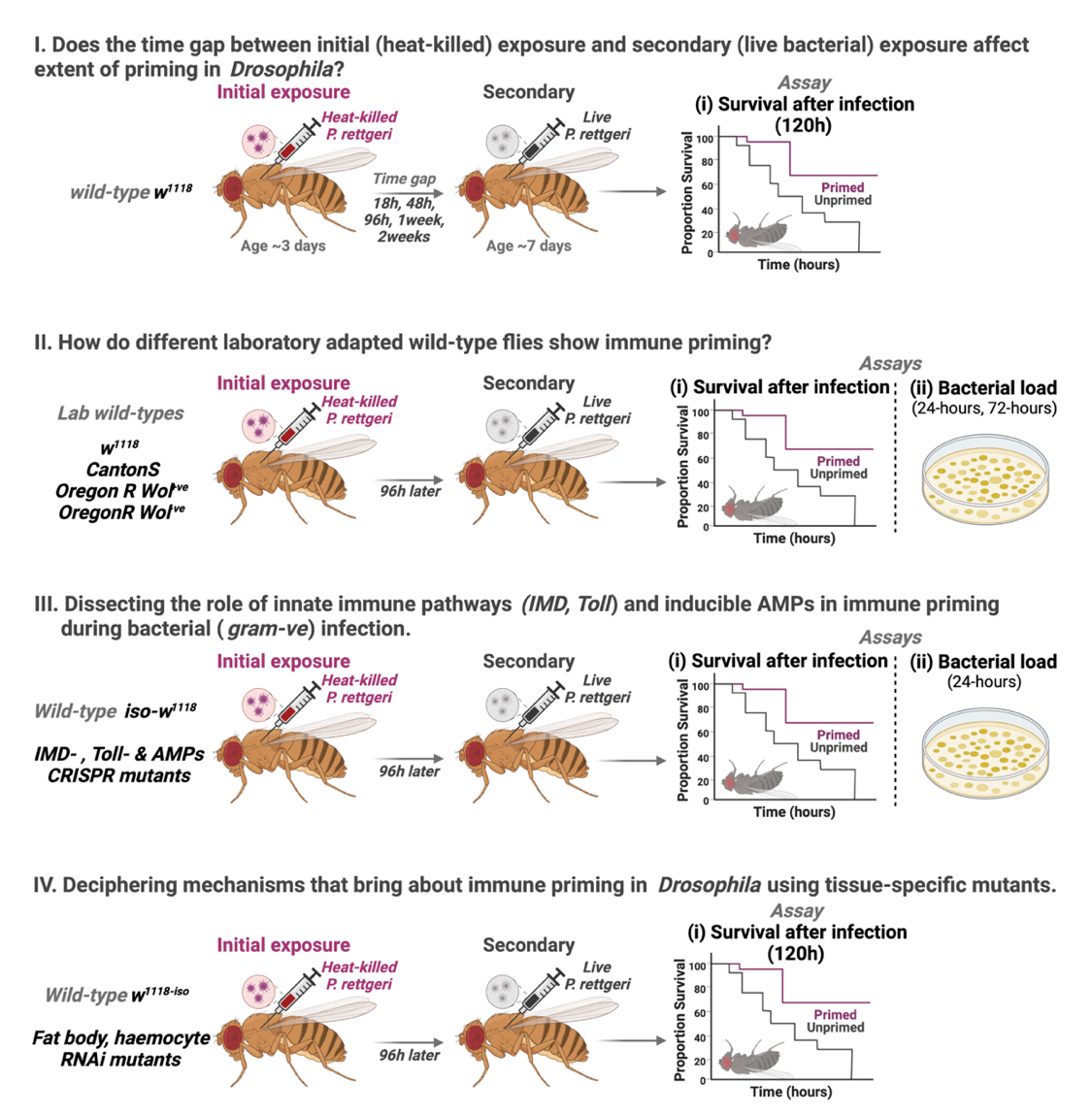

Figure S2: Experimental design to measure epidemiological components following systemic (OD_600_=1) and oral (OD_600_=25) priming and infection with initial heat-killed exposure followed by live *P. rettgeri* in male and female wildtype w^1118^ flies. The assays include **(I).** survival following different infection routes **(II).** internal bacterial load (III). behavioural components of pathogen exposure such as sleep and awake activity **(IV).** bacteria shedding and **(V).** transmission. n=6-7 vials of 8-12 flies in each vial, for each treatment and sex combination.

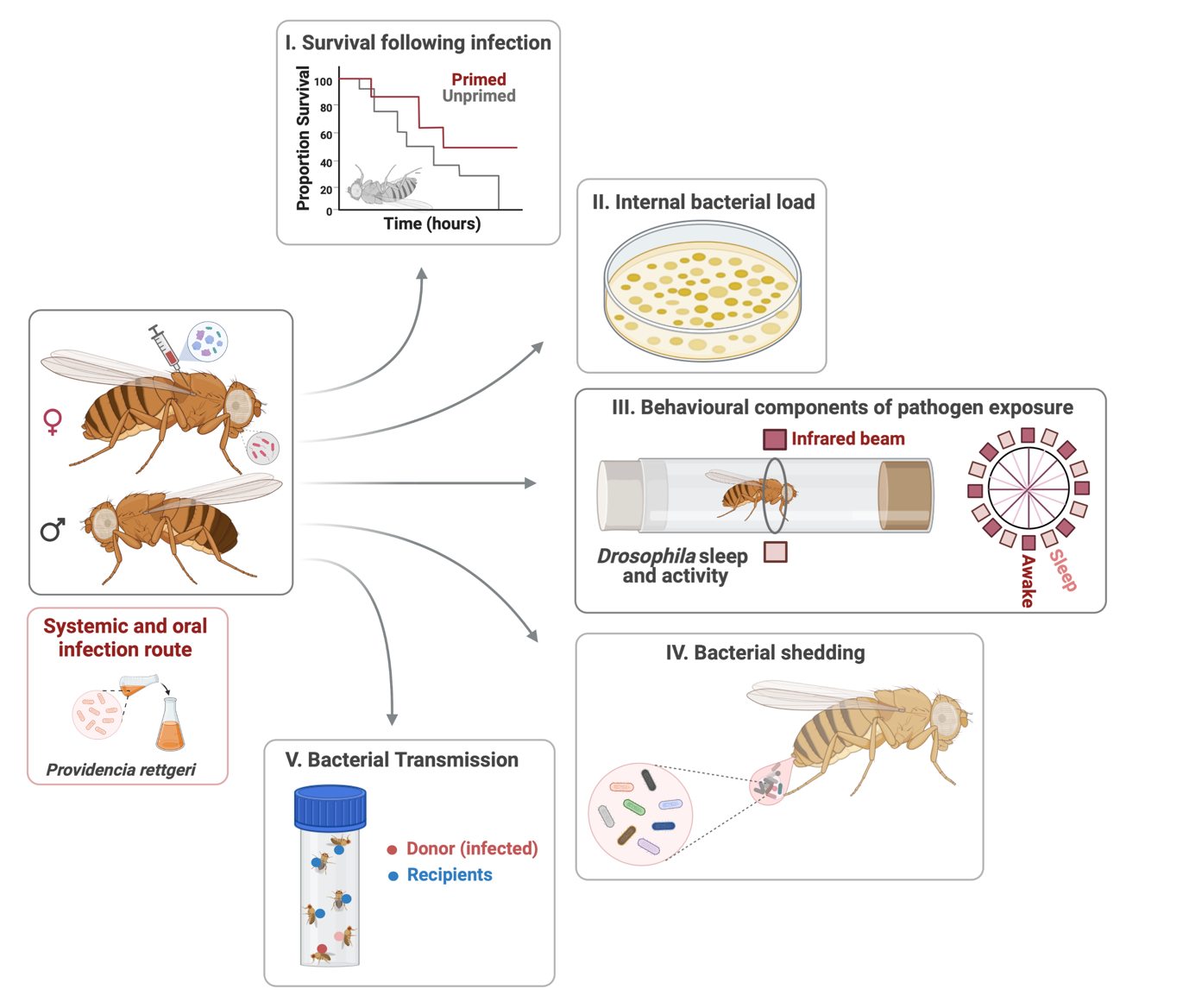

Figure S3. Survival curves of w1118 and iso-w1118 (Drosdel) flies after initial heat-killed exposure and followed by live *P. rettgeri* infection with OD_600_ = 0.1. As another control, we infected both w^1118^ and iso-w^1118^ control because the CRISPR/cas9 AMP mutants we used were on the *iso-w^1118^* background, so we wanted to confirm that any changes in priming were not due to the background of the mutants, as opposed to the mutations. We found that the differences between *w^1118^* and *iso-w^1118^* (primed and unprimed treatments) were not significantly different.

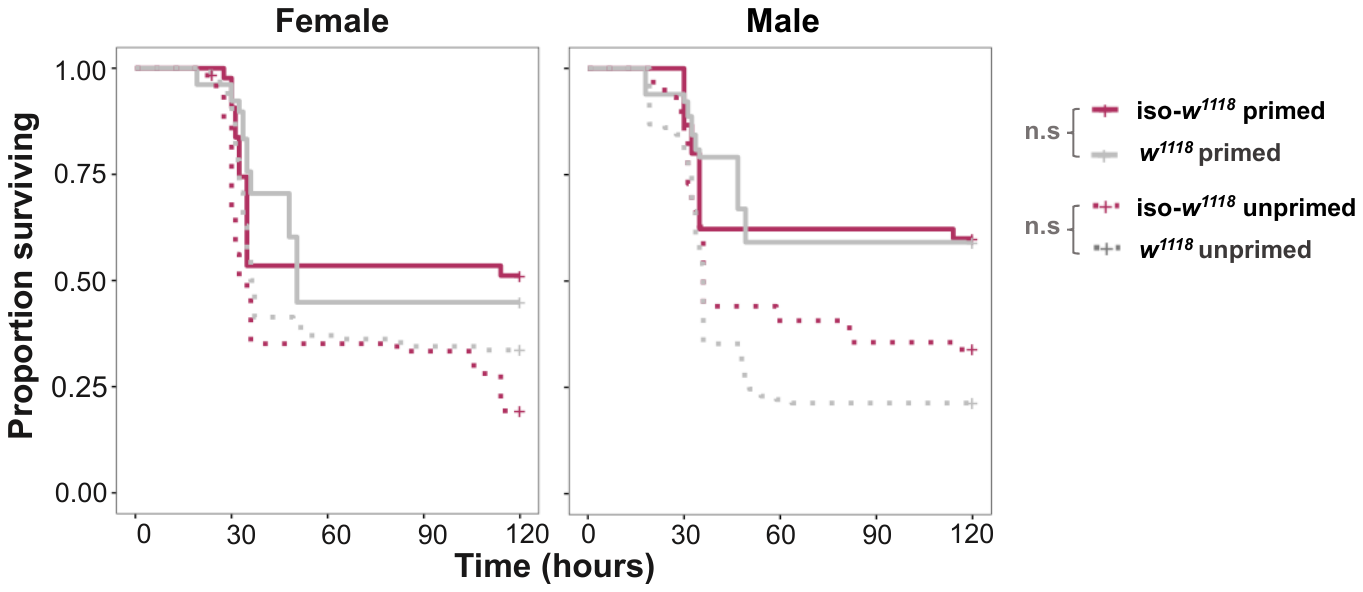

### Supplementary tables

Table-S1. Summary of mixed effects Cox model, fitting the model to estimate time-delayed priming response in control w1118 male and female flies. We used data from individuals exposed to live bacterial after different time intervals of initial heat-killed *P. rettgeri* exposure. We specified the model as: survival ~ treatment x sex x timepoint (1|vial/block), with treatment and sex as fixed effects, and vials nested within each block and as a random effect. The table shows model output (ANOVA) for priming in control flies.

| **Timepoint/sex** | **Source** | **loglik** | **χ2** | **Df** | **P** |
| --- | --- | --- | --- | --- | --- |
| ***18-hours*** *post*  *Priming overall* | Treatment  Sex  Sex × Treatment | -801.96  -790.74 | 15.92  22.44 | 1  1 | **<0.001**  **<0.001** |
|  |  | - 781.47 | 18.54 | 1 | **<0.001** |
|  | *Random effects*  *Vials/block* | *Std Dev* |  |  |  |
|  |  | *0.08* |  |  |  |
| *Female* | Treatment | -433.36 | 0.64 | 1 | 0.42 |
|  | *Random effects*  *Vials/block* | *Std Dev*  *0.18* |  |  |  |
| *Male* | Treatment | -247.71 | 32.37 | 1 | **<0.001** |
|  | *Random effects* | *Std Dev* |  |  |  |
|  | *Vials/block* | *0.009* |  |  |  |
| ***48-hours*** *post*  *priming* | Treatment  Sex  Sex × Treatment | -1050.2  - 1045.7 | 25.73  8.94 | 1  1 | **<0.001**  **0.002** |
| *overall* |  | -1045.7 | 0.08 | 1 | 0.76 |
|  | *Random effects*  *Vials/block* | *Std Dev* |  |  |  |
|  |  | *0.004* |  |  |  |
| *Female* | Treatment | -315.12 | 21.15 | 1 | **<0.001** |
|  | *Random effects*  *Vials/block* | *Std Dev*  *0.23* |  |  |  |
| *Male* | Treatment | -239.08 | 15.63 | 1 | **<0.001** |
|  | *Random effects* | *Std Dev* |  |  |  |
|  | *Vials/block* | *0.15* |  |  |  |
| ***96-hours*** *post*  *Priming* | Treatment  Sex  Sex × Treatment | -647.58  -647.51 | 36.35  0.14 | 1  1 | **<0.001**  0.70 |
| *overall* |  | -647.50 | 0.006 | 1 | 0.93 |
|  | *Random effects* | *Std Dev* |  |  |  |
|  | *Vials/block* | 0.15 |  |  |  |
| *Female* | Treatment | -511.02 | 14.63 | 1 | **<0.001** |
|  | *Random effects* | *Std Dev* |  |  |  |
|  | *Vials/block* | *0.009* |  |  |  |
| *Male* | Treatment | -399.48 | 12.69 | 1 | **<0.001** |
|  | *Random effects*  *Vials/block* | *Std Dev* |  |  |  |
|  |  | *0.008* |  |  |  |
| ***168-hours*** *post*  *Priming*  *overall* | Treatment  Sex  Sex × Treatment | -949.67  -948.59  -946.20 | 11.712  2.1658  4.7705 | 1  1  1 | **0.0006**  0.14  **0.02** |
|  | *Random effects* | *Std Dev* |  |  |  |
|  | *Vials/block* | *0.009* |  |  |  |
| *Female* | Treatment | -448.02 | 1.1289 | 1 | 0.28 |
|  | *Random effects* | *Std Dev* |  |  |  |
|  | *Vials/block* | *0.009* |  |  |  |
| *Male* | Treatment | -371.76 | 13.381 | 1 | **<0.001** |
|  | *Random effects* | *Std Dev* |  |  |  |
|  | *Vials/block* | *0.009* |  |  |  |
| ***336-hours*** *post*  *Priming*  *overall* | Treatment  Sex  Sex × Treatment | -1035.0  -1034.9  -1033.5 | 5.3983  0.1854  2.7814 | 1  1  1 | **0.02**  0.66  0.09 |
|  | *Random effects* | *Std Dev* |  |  |  |
|  | *Vials/block* | *0.004* |  |  |  |
| *Female* | Treatment | -445.91 | 0.2168 | 1 | 0.64 |
|  | *Random effects* | *Std Dev* |  |  |  |
|  | *Vials/block* | *0.01* |  |  |  |
| *Male* | Treatment | -448.08 | 8.345 | 1 | **0.003** |
|  | *Random effects* | *Std Dev* |  |  |  |
|  | *Vials/block* | *0.008* |  |  |  |

Table-S2. Summary of mixed effects Cox model, fitting the model to estimate priming response in different laboratory control w1118 male and female flies. We used data from the unprimed-infected and the primed-infected treatments and specified the model as: survival ~ treatment x sex x (1|vial/block), with treatment and sex as fixed effects, and vials nested within each block and as a random effect. The table shows model output (ANOVA) for priming in control flies.

| **Fly strain** | **Source** | **loglik** | **χ2** | **Df** | **P** |
| --- | --- | --- | --- | --- | --- |
| *w^1118^* | Treatment  Sex  Sex × Treatment | -647.58  -647.51 | 36.35  0.146 | 1  1 | **<0.001**  0.70 |
|  |  | -647.50 | 0.007 | 1 | 0.93 |
|  | *Random effects*  *Vials/block* | *Std Dev* |  |  |  |
|  |  | *0.15* |  |  |  |
| *Canton-S* | Treatment  Sex  Sex × Treatment | -633.00  -619.47 | 11.56  27.04 | 1  1 | **<0.001**  **<0.001** |
|  |  | -618.39 | 2.174 | 1 | 0.14 |
|  | *Random effects*  *Vials/block* | *Std Dev* |  |  |  |
|  |  | *0.008* |  |  |  |
| *OreR^Wol+^* | Treatment  Sex  Sex × Treatment | -436.34  -436.04 | 0.455  0.607 | 1  1 | 0.49  0.43 |
|  |  | -432.80 | 6.482 | 1 | **0.01** |
|  | *Random effects*  *Vials/block* | *Std Dev* |  |  |  |
|  |  | *0.17* |  |  |  |
| *OreR^Wol-^* | Treatment  Sex  Sex × Treatment | -610.99  -609.91 | 26.18  2.171 | 1  1 | **<0.001**  0.14 |
|  |  | -604.13 | 11.54 | 1 | **<0.001** |
|  | *Random effects*  *Vials/block* | *Std Dev* |  |  |  |
|  |  | *0.02* |  |  |  |

Table-S3. Summary of mixed effects Cox model, fitting the model to estimate the impact Wolbachia on immune priming response using genetic background OreR male and female flies. We used data from the unprimed-infected and the primed-infected treatments and specified the model as: survival ~ treatment x sex x *Wolbachia* status (1|vial/block), with treatment and sex as fixed effects, and vials nested within each block and as a random effect. The table shows model output (ANOVA) for priming in different control/genetic background flies.

A. **Impact of *Wolbachia***

| **Fly strain** | **Source** | **loglik** | **χ2** | **Df** | **P** |
| --- | --- | --- | --- | --- | --- |
| *OreR* | Treatment  Sex  *Wolbachia* status  Sex × Treatment  Treatment x *Wol* status  Sex x *Wol* status  Sex x Treatment x *Wol* status | -1213.1  -1210.9  -1197.4 | 12.88  4.31  27.06 | 1  1  1 | **<0.001**  **0.037**  **<0.001** |
|  |  | -1189.5  -1183.8  -1183.4  -1183.4 | 15.84  11.36  0.69  0.06 | 1  1  1  1 | **<0.001**  **<0.001**  0.40  0.79 |
|  | *Random effects*  *Vials/block* | *Std Dev* |  |  |  |
|  |  | *0.004* |  |  |  |

B. ***Wolbachia* infection on males and females separately**

| **Sex** | **Source** | **loglik** | **χ2** | **Df** | **P** |
| --- | --- | --- | --- | --- | --- |
| *Female* | Treatment  *Wol* status  Treatment x *Wol* status | -514.76  -506.29  -503.72 | 0.10  16.94  5.13 | 1  1  1 | 0.74  **<0.001**  **0.02** |
|  | *Random effects* | *Std Dev* |  |  |  |
|  | *Vials/block* | 0.008 |  |  |  |
| *Male* | Treatment  *Wol* status  Treatment x *Wol* status | -535.18  -529.14  -525.88 | 23.90  12.07  6.52 | 1  1  1 | **<0.001**  **<0.001**  **0.01** |
|  | *Random effects* | *Std Dev* |  |  |  |
|  | *Vials/block* | 0.009 |  |  |  |

Table-S4. A. Summary of mixed effects Cox model, fitting the model to estimate priming response in male and w1118 flies primed and challenged with homologous and heterologous combination of bacteria. We used data from the unprimed-infected, primed-infected with same bacterial species, primed-infected with heterologous treatments and specified the model as: survival ~ Treatment, with treatment as fixed effect. The table shows model output (ANOVA).

| **A. Source** | **df** | **χ2** | **p** |
| --- | --- | --- | --- |
| Treatment | 3 | 29.51 | <0.001 |

B. Summary of the estimated hazard ratios for male *w^1118^* flies (homologous and heterologous priming-challenge combinations), including their lower and upper limits of the 95% confidence interval, calculated from survival curves (Fig. 3A, n = 7 vials/treatment combinations). A greater hazard ratio (>1) indicates higher susceptibility to infection.

| **B. Combinations** | **treatment** |  | **Hazard ratio** | **p** | **Lower 95%** | **Upper 95%** |
| --- | --- | --- | --- | --- | --- | --- |
| Homologous | *Pr-Pr vs* | *PBS-Pr* | 0.58 | 0.01 | 0.38 | 0.88 |
| Heterologous | *Pr-Pb vs* | *PBS-Pr* | 1.02 | 0.88 | 0.7 | 1.49 |
|  | *Pr-Pe vs* | *PBS-Pr* | 1.71 | 0.004 | 1.18 | 2.47 |
|  | *Pr-Pb vs* | *Pr-Pr* | 1.75 | 0.005 | 1.18 | 2.6 |
|  | *Pr-Pe vs* | *Pr-Pr* | 2.92 | 0.001 | 1.97 | 4.32 |

Table-S5. Summary of mixed effects Cox model, fitting the model to estimate priming response in male and female control w1118 flies. We used data from the unprimed-infected and the primed-infected treatments and specified the model as: survival ~ Treatment x sex x (1|vial/block), with treatment and sex as fixed effects, and vials within a block as a random effect. The table shows model output (ANOVA).

| **Fly strain** | **Source** | **loglik** | **χ2** | **Df** | **P** |
| --- | --- | --- | --- | --- | --- |
| *w^1118^* | Sex  Treatment  Sex × Treatment | -5144.0  -5163.8 | 39.56  0.351 | 1  1 | **<0.001**  **<0.001** |
|  |  | -5143.5 | 0.921 | 1 | 0.34 |
|  | *Random effects*  *Vials/block* | *Std Dev* |  |  |  |
|  |  | *0.453* |  |  |  |

***Table-S6.*** Summary of log_10_ transformed bacterial load data after 0.2 OD *P. rettgeri* infection, analysed using a non-parametric test for ANOVA (Kruskal-Wallis test) by fitting ‘treatment’, as fixed-effects for female and male control *w^1118^.*

| **Time** | **Sex** | **Source** | **Chi Sq.** | **Df** | **P** |
| --- | --- | --- | --- | --- | --- |
| ***24-hours*** *following infection* | *Female*  *Male* | Treatment  Treatment | 3.2495  8.3974 | 1  1 | 0.07  **0.003** |
| ***72-hours*** *following infection* | *Female*  *Male* | Treatment  Treatment | 0.2102  1.8293 | 1  1 | 0.64  0.17 |

Table-S7: Summary of Cox prop-hazard model, for female and male flies of wild type w1118. We used data from 3 to 7-day adult male and females infected with OD_600_ = 0.75 dose for systemic and dose OD_600_ = 25 for oral priming and infection with *P. rettegeri*. We specified the models as survival ~ sex x treatment with ‘treatment’, and ‘sex’ as fixed effects. Table shows model output (ANOVA) for survival post-infection for male and female *w^1118^* flies.

| ***Response*** | ***Predictor*** | ***df*** | ***Chi sq*** | ***p*** |
| --- | --- | --- | --- | --- |
| **Systemic Infection route** | Sex | 1 | 1.383 | 0.23 |
|  | Treatment | 2 | 100.7 | **<0.001** |
|  | Sex x Treatment | 2 | 3.103 | 0.21 |
| **Oral Infection route** | Sex | 1 | 0.728 | 0.39 |
|  | Treatment | 2 | 85.76 | **<0.001** |
|  | Sex x Treatment | 2 | 3.421 | 0.180 |

Table-S8: Summary of log transformed bacterial load data after OD_600_=0.75 systemic and OD_600_=25 oral *P. rettgeri* systemic infection for male and female *w^1118^* flies analysed using a non-parametric Wilcoxon (Kruskal-Wallis) test by fitting ‘Sex’ as categorical fixed-effects after 24-hours post systemic and oral priming and infection.

| ***Response*** | ***Sex*** | ***Predictor*** | ***Chi sq*** | ***df*** | ***p*** |
| --- | --- | --- | --- | --- | --- |
| ***Systemic route*** | Female | Treatment | 0.403 | 1 | 0.52 |
|  | Male | Treatment | 6.090 | 1 | **0.01** |
| ***Oral route*** | Female | Treatment | 0.255 | 1 | 0.61 |
|  | Male | Treatment | 3.870 | 1 | **0.04** |

Table-S9: Model outputs for statistical test (GLM) performed on host activity data, that is, locomotor activity, sleep patterns and average awake activity in males and females of *w^1118^* during systemic and oral priming and infection respectively.

| ***Infection*** | ***Response*** | ***Predictor*** | ***df*** | ***F ratio*** | ***p*** |
| --- | --- | --- | --- | --- | --- |
| ***Oral*** | *Total activity* | Sex | 1 | 16.82 | **<0.001** |
|  |  | Treatment | 2 | 0.711 | 0.49 |
|  |  | Sex x Treat | 2 | 0.020 | 0.97 |
|  | *Awake activity* | Sex | 1 | 40.86 | **<0.001** |
|  |  | Treatment | 2 | 0.269 | 0.76 |
|  |  | Sex x Treat | 2 | 0.112 | 0.89 |
|  | *Time asleep* | Sex | 1 | 40.86 | **<0.001** |
|  |  | Treatment | 2 | 0.269 | 0.76 |
|  |  | Sex x Treat | 2 | 0.112 | 0.89 |
| ***Systemic*** | *Total activity* | Sex | 1 | 0.612 | 0.43 |
|  |  | Treatment | 2 | 19.18 | **<0.001** |
|  |  | Sex x Treat | 2 | 0.294 | 0.74 |
|  | *Awake activity* | Sex | 1 | 0.555 | 0.45 |
|  |  | Treatment | 2 | 41.03 | **<0.001** |
|  |  | Sex x Treat | 2 | 0.238 | 0.78 |
|  | *Time asleep* | Sex | 1 | 0.555 | 0.45 |
|  |  | Treatment | 2 | 41.03 | **<0.001** |
|  |  | Sex x Treat | 2 | 0.238 | 0.78 |

Table-S10: Summary of non-parametric Wilcoxon (Kruskal-Wallis) test for oral bacterial shedding (log transformed bacterial load) after 4-hours of oral priming and infection with *P. rettgeri* OD_600_=25. Bacterial transmission (percentage of recipient flies with measurable bacterial load) after 4-hours of exposure with infection (donor) *w^1118^* after oral priming and infection.

| ***Response*** | ***Sex*** | ***Predictor*** | ***Chi sq*** | ***df*** | ***p*** |
| --- | --- | --- | --- | --- | --- |
| ***Bacterial Shedding*** | Female | Treatment | 0.003 | 1 | 0.95 |
|  | Male | Treatment | 17.29 | 1 | **<0.001** |
| ***Transmission*** | Female | Treatment | 0.187 | 1 | 0.66 |
|  | Male | Treatment | 5.444 | 1 | **0.019** |

Table-S11. Summary of mixed effects Cox model, fitting the model to estimate priming response in male and female control w1118, IMD and Toll transgenic flies. We used data from the unprimed-infected and the primed-infected treatments and specified the model as: survival ~ Treatment x sex x (1|vial/block), with treatment and sex as fixed effects, and vials within a block as a random effect for each fly line. The table shows model output (ANOVA).

| **Fly strain** | **Source** | **loglik** | **χ2** | **Df** | **P** |
| --- | --- | --- | --- | --- | --- |
| *iso w^1118^* | Sex  Treatment  Sex × Treatment | -1597.3  -1594.0 | 6.734  47.70 | 1  1 | **0.009**  **<0.001** |
|  |  | -1594.0 | 0.000 | 1 | 0.994 |
|  | *Random effects*  *Vials/block* | *Std Dev* |  |  |  |
|  |  | *4.1* |  |  |  |
| *Rel^E20^* | Sex  Treatment  Sex × Treatment | -1497.2  -1497.3 | 0.048  3.56 | 1  1 | 0.82  0.058 |
|  |  | -1497.1 | 0.20 | 1 | 0.65 |
|  | *Random effects*  *Vials/block* | *Std Dev* |  |  |  |
|  |  | *0.26* |  |  |  |
| *𝝙 AMPs* | Sex  Treatment  Sex × Treatment | -1605.6  -1605.7 | 0.263  1.920 | 1  1 | 0.60  0.16 |
|  |  | -1604.9 | 1.451 | 1 | 0.22 |
|  | *Random effects*  *Vials/block* | *Std Dev* |  |  |  |
|  |  | *0.21* |  |  |  |
| *Group-B* | Sex  Treatment  Sex × Treatment | -1537.3  -1538.8 | 2.950  0.299 | 1  1 | 0.08  0.58 |
|  |  | -1536.9 | 0.771 | 1 | 0.37 |
|  | *Random effects*  *Vials/block* | *Std Dev* |  |  |  |
|  |  | *0.02* |  |  |  |
| *Dpt* | Sex  Treatment  Sex × Treatment | -3386.1  -3398.4 | 24.49  3.157 | 1  1 | **<0.001**  0.08 |
|  |  | -3386.1 | 0.105 | 1 | 0.74 |
|  | *Random effects*  *Vials/block* | *Std Dev* |  |  |  |
|  |  | *0.03* |  |  |  |
| *𝝙 AMPs^+Dpt^* | Sex  Treatment  Sex × Treatment | -2596.1  -2613.2 | 34.22 23.92 | 1  1 | **<0.001**  **<0.001** |
|  |  | -2580.5 | 31.10 | 1 | **<0.001** |
|  | *Random effects*  *Vials/block* | *Std Dev* |  |  |  |
|  |  | *3.73* |  |  |  |
| *Spz* | Sex  Treatment  Sex × Treatment | -1755.6  -1763.0 | 14.86 21.61 | 1  1 | **<0.001**  **<0.001** |
|  |  | -1755.4 | 0.298 | 1 | 0.58 |
|  | *Random effects*  *Vials/block* | *Std Dev* |  |  |  |
|  |  | *0.08* |  |  |  |

Table-S12. Summary of log10 transformed bacterial load data in AMP deletion lines after 0.2 OD *P. rettgeri* infection, analysed using non-parametric ANOVA (K-W test) by fitting ‘treatment’ (i.e., primed and unprimed) as categorical fixed-effects for male and females of each fly lines (control *w^1118^* and transgenic flies).

| **Fly line** | **Sex** | **Chi Sq.** | **Df** | **P** |
| --- | --- | --- | --- | --- |
| *w^1118^* | *Female* | 1.1165 | 1 | 0.29 |
|  | *Male* | 7.5172 | 1 | **0.006** |
| *Rel^E20^* | *Female*  *Male* | 4.0368  0.2516 | 1  1 | **0.044**  0.61 |
| *Spz* | *Female*  *Male* | 0.0152  0.0050 | 1  1 | 0.90  0.94 |
| *𝝙 AMPs* | *Female*  *Male* | 4.9868  0.3014 | 1  1 | **0.025**  0.58 |
| *Group-B* | *Female*  *Male* | 0.5612  3.5402 | 1  1 | 0.45  0.059 |
| *Dpt* | *Female*  *Male* | 0.2848  0.0020 | 1  1 | 0.59  0.96 |
| *𝝙 AMPs^+Dpt^* | *Female*  *Male* | 0.1589  11.2941 | 1  1 | 0.69  **0.0008** |

Table-S13. Summary of mixed effects Cox prop-hazard in tissue-speciifc Dpt knockdown lines, fitting the model to estimate strength of priming response in male and female control w^1118^ and UAS^-RNAi^ tissue-specific mutants. We used data from the unprimed-infected and the primed-infected treatments and specified the model as: survival ~ Treatment x sex, with treatment and sex as fixed effects for each fly line. The table shows model output (ANOVA).

| **Fly strain** | **Source** | **χ2** | **Df** | **P** |
| --- | --- | --- | --- | --- |
| *w^1118-iso^* | Sex  Treatment  Sex × Treatment | 2.306 | 1 | 0.12 |
|  |  | 11.95  3.226 | 1  1 | **0.005**  0.075 |
| *FB>DptB* | Sex  Treatment  Sex × Treatment | 1.996 | 1 | 0.15 |
|  |  | 0.977  0.068 | 1  1 | 0.32  0.79 |
| *HH>DptB* | Sex  Treatment  Sex × Treatment | 3.609 | 1 | 0.057 |
|  |  | 17.06  0.602 | 1  1 | **<0.001**  0.43 |

Table-S14. Summary of log10 transformed Dpt gene expression data in control w^1118^ flies after 0.2 OD *P. rettgeri* priming and challenge, analysed using ANOVA by fitting ‘treatment’ and ‘sex’ as categorical fixed-effects.

| **Condition** | **Source** | **F value** | **Df** | **P** |
| --- | --- | --- | --- | --- |
| Before secondary exposure (***Dpt***) | Sex  Treatment  Sex × Treatment | 36.96  5.295  0.297 | 1  1  1 | **<0.001**  **<0.001**  0.13 |
| After secondary exposure (***Dpt***) | Sex  Treatment  Sex × Treatment | 0.487  29.48  2.465 | 1  1  1 | 0.49  **<0.001**  0.13 |
| ***AttC*** and ***Dro*** AMP genes expression after secondary exposure | AMP genes  Treatment  Timepoint  Treatment x genes | 98.53  1.704  29.51  0.725 | 1  1  1  1 | **<0.001**  0.20  **<0.001**  0.40 |
|  | Treatment x timepoint | 0.028 | 1 | 0.86 |
|  | Gene x timepoint | 60.07 | 1 | **<0.001** |
|  | Treatment x gene x timepoint | 0.323 | 1 | 0.57 |

Table-S15. Summary of mixed effects Cox model, fitting the model to estimate priming response in male and female PGRP mutant flies. We used data from the unprimed-infected and the primed-infected treatments and specified the model as: survival ~ Treatment x sex x (1|vial), with treatment and sex as fixed effects, and vials within a block as a random effect for each fly line. The table shows model output (ANOVA).

| **Fly strain** | **Source** | **loglik** | **χ2** | **Df** | **P** |
| --- | --- | --- | --- | --- | --- |
| *PGRP-LB* | Treatment  Sex  Sex × Treatment | -818.79  -818.13 | -0.0048  1.3085 | 1  1 | 1.00  0.25 |
|  |  | -817.98 | 0.3047 | 1 | 0.58 |
|  | *Random effects*  *Vials/block* | *Std Dev* |  |  |  |
|  |  | *9.011949e-03* |  |  |  |
| *PGRP-LC* | Treatment  Sex  Sex × Treatment | -891.65  -891.65 | 0.0189  0.0057 | 1  1 | 0.89  0.96 |
|  |  | -891.64 | 0.0068 | 1 | 0.93 |
|  | *Random effects*  *Vials/block* | *Std Dev* |  |  |  |
|  |  | *9.281555e-03* |  |  |  |
| *PGRP-LE* | Treatment  Sex  Sex × Treatment | -928.63  -928.31 | 0.2609  0.6362 | 1  1 | 0.60  0.42 |
|  |  | -928.31 | 0.0036 | 1 | 0.95 |
|  | *Random effects*  *Vials/block* | *Std Dev* |  |  |  |
|  |  | *0.17* |  |  |  |

Table-S16. Summary of log10 transformed bacterial load data after 0.2 OD P. rettgeri infection, analysed using non-parametric ANOVA (Kruskal-Wallis test) by fitting ‘treatment’ (i.e., primed and unprimed) as categorical fixed-effects for male and females of PGRP mutants.

| **Fly line** | **Sex** | **Chi Sq.** | **Df** | **P** |
| --- | --- | --- | --- | --- |
| *PGRP-LB* | *Female* | 0.4300 | 1 | 0.51 |
|  | *Male* | 0.3295 | 1 | 0.56 |
| *PGRP-LC* | *Female*  *Male* | 0.6063  0.2020 | 1  1 | 0.43  0.65 |
| *PGRP-LE* | *Female*  *Male* | 0.0643  0.0813 | 1  1 | 0.79  0.77 |

Table-S17. Summary of log10 transformed Dpt gene expression data in flies with different PGRP deletion transgenic flies after 0.2 OD *P. rettgeri* priming and challenge, analysed using ANOVA by fitting ‘fly line’ (*PGRP-LB, -LC & -LE*) ‘treatment’ (*primed, unprimed*) and ‘sex’ as categorical fixed-effects

| **Source** | **df** | **Sum of Sq.** | **F ratio** | **p** |
| --- | --- | --- | --- | --- |
| Fly line | 2 | 596443.9 | 582.0 | **<.0001** |
| Sex | 1 | 2981.64 | 5.819 | **0.02** |
| Fly line x sex | 2 | 5623.86 | 5.487 | **0.007** |
| Treatment | 1 | 783.67 | 1.529 | 0.22 |
| Fly line x treatment | 2 | 1754.66 | 1.712 | 0.19 |
| Sex x treatment | 1 | 267.07 | 0.521 | 0.47 |
| Fly line sex x treat | 2 | 567.95 | 0.554 | 0.57 |
